# Multiscale and integrative single-cell Hi-C analysis with Higashi

**DOI:** 10.1101/2020.12.13.422537

**Authors:** Ruochi Zhang, Tianming Zhou, Jian Ma

## Abstract

The advent of single-cell Hi-C (scHi-C) technologies offers an unprecedented opportunity to unveil cell-to-cell variability of 3D genome organization. However, the development of computational methods that can effectively enhance scHi-C data quality and extract 3D genome features in single cells remains a major challenge. Here, we report Higashi, a new algorithm that achieves state-of-the-art analysis of scHi-C data based on hypergraph representation learning. Extensive evaluations demonstrate that Higashi significantly outperforms existing methods for embedding and imputation of scHi-C data. Higashi is uniquely able to identify multiscale 3D genome features (such as compartmentalization and TAD-like domain boundaries) in single cells, allowing markedly refined delineation of cell-to-cell variability of 3D genome features. By applying to a scHi-C dataset from human prefrontal cortex, Higashi reveals complex cell types as well as new connections between 3D genome features and cell type-specific gene regulation. Higashi provides an end-to-end solution to scHi-C data analysis and is applicable to studying single-cell 3D genomes in a wide range of biological contexts.

## Introduction

The rapid development of whole-genome mapping methods such as Hi-C (Lieberman-Aiden et al., 2009) for probing the three-dimensional (3D) genome organization inside the nucleus has revealed multiscale higher-order chromatin structures (Kempfer and Pombo, 2020), including A/B compartments (Lieberman-Aiden et al., 2009), more refined nuclear compartmentalization (Rao et al., 2014; Xiong and Ma, 2019; Wang et al., 2021), topologically-associating domains (TADs) (Dixon et al., 2012; Nora et al., 2012), and chromatin loops (Rao et al., 2014). These 3D genome features in different scales are interconnected with vital genome functions such as gene transcription and DNA replication (Dekker et al., 2017; Marchal et al., 2019), yet the variation of 3D genome structures and its functional implication in single cells remain mostly unclear (Misteli, 2020). The emerging single-cell Hi-C (scHi-C) technologies have enabled genomic mapping of 3D chromatin structures in individual cells (Nagano et al., 2013; Stevens et al., 2017; Flyamer et al., 2017; Ramani et al., 2017; Nagano et al., 2017; Tan et al., 2018) and, more recently, joint profiling of chromosome conformation with other epigenomic features (Lee et al., 2019; Li et al., 2019). These exciting scHi-C assays have the potential to comprehensively reveal fundamental genome structure and function connections at single-cell resolution in a wide range of biological contexts.

However, computational methods that can fully leverage the sparse scHi-C data to analyze the cell-to-cell variability of 3D genome features are significantly lacking. To account for the sparseness of scHi-C data, methods have been developed for embedding the datasets (Liu et al., 2018; Kim et al., 2020) and the imputation of the contact maps (Zhou et al., 2019). However, the current state-of-the-art imputation methods based on “random walk with restart” such as scHiCluster (Zhou et al., 2019) have much room for improvement for more reliable single-cell 3D genome analysis. Current imputation methods also require storage and calculation on dense matrices with the size of the contact maps in memory, which is impractical when analyzing scHi-C data at relatively high resolutions. It also remains unclear how to reliably compare TAD-like domain boundaries and A/B compartments across single cells to analyze their cell-to-cell variability and functional connections. Therefore, new algorithms are urgently needed to fill these important gaps.

Here, we report Higashi, a new computational method for multiscale and integrative single-cell Hi-C analysis using hypergraph representation learning. Using the embeddings and the imputed scHi-C contact maps produced by Higashi, we identified cell-to-cell variability of A/B compartment scores and TAD-like domain boundaries that are functionally important. Application to a recent scHi-C dataset of human prefrontal cortex demonstrated the unique ability of Higashi to reveal cell type-specific 3D genome features in complex tissues. As a new and the most systematic method to date, Higashi enables much improved analysis of scHi-C data with the potential to shed new light on the dynamics of 3D genome structures and their functional implications in different biological processes.

## Results

### Overview of Higashi

The key algorithmic design of Higashi is to transform the scHi-C data into a hypergraph (Fig. 1a). Such transformation preserves the single-cell resolution and the 3D genome features from the scHi-C contact maps. Specifically, the process of embedding the scHi-C data is now equivalent to learning node embeddings of the hypergraph, while imputing the scHi-C contact maps becomes predicting missing hyperedges within the hypergraph. In Higashi, we leverage our recently developed Hyper-SAGNN architecture (Zhang et al., 2020), which is a generic hypergraph representation learning framework, with substantial new development specifically for scHi-C analysis (see Methods).

**Figure 1:**
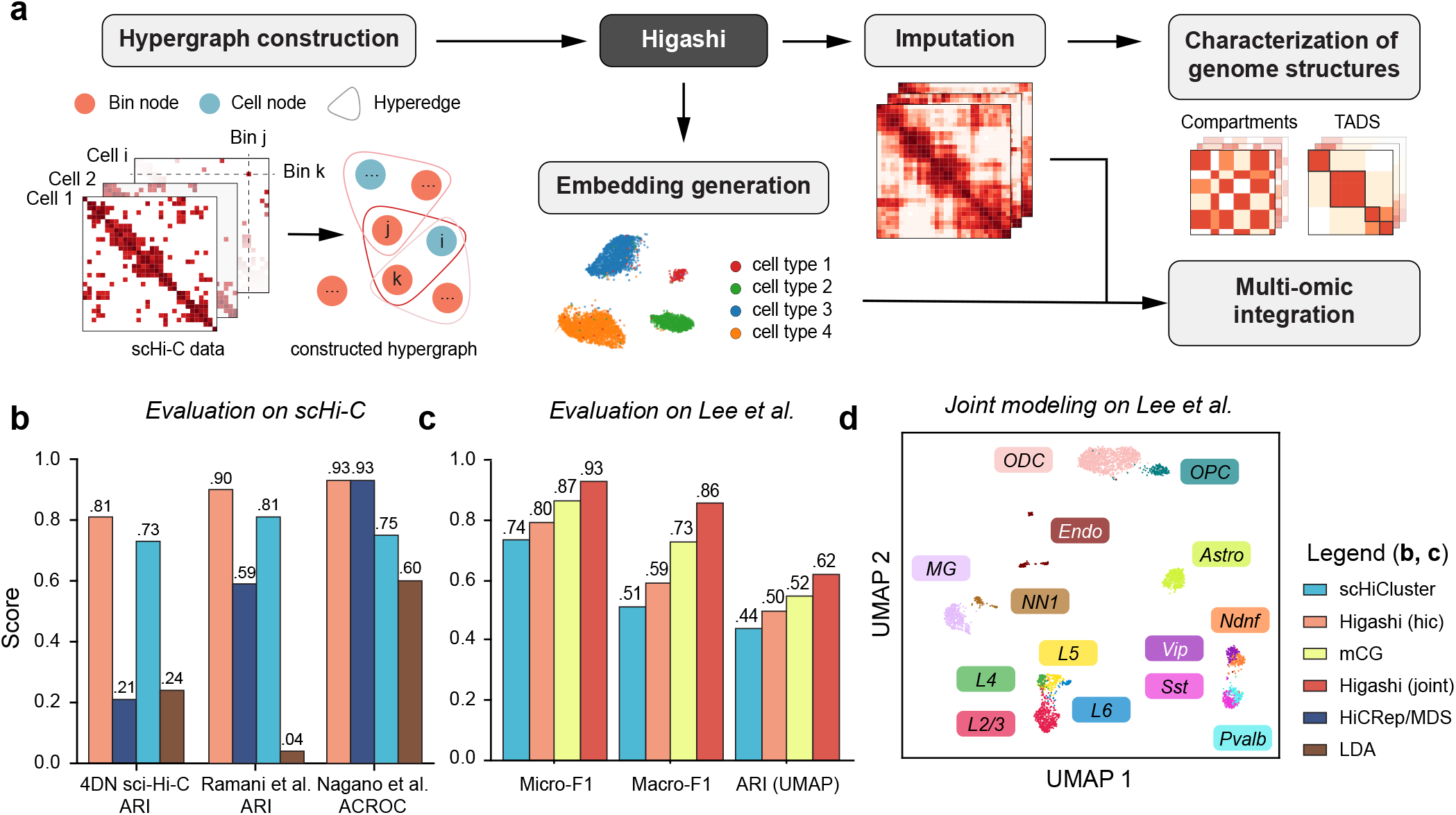
Overall design of Higashi and the evaluation of Higashi embeddings on real data. **a**. Overview of the Higashi framework for scHi-C analysis. The input scHi-C dataset is transformed into a hypergraph where each hyperedge connects one cell node and two bin nodes. A hypergraph neural network is trained to capture high-order interaction patterns within the constructed hypergraph. The trained neural network is able to generate embeddings for scHi-C data and impute the sparse scHi-C contact maps. The imputed contact maps and the embeddings allow detailed characterization of multiscale 3D genome features and also multiomic integrative analysis. **b**. Quantitative evaluation of Higashi on the three public scHi-C datasets by comparing to HiCRep/MDS (Liu et al., 2018), scHiCluster (Zhou et al., 2019), and LDA (Kim et al., 2020). The performances are measured by Adjusted Rand Index (ARI), and also ACROC scores from the unsupervised cell type identification tasks. See also Fig. S2. **c**. Quantitative evaluation of different embeddings of the sn-m3C-seq data (Lee et al., 2019) using Micro-F1, Macro-F1, and Adjusted Rand Index (ARI) scores. **d**. UMAP visualization of the Higashi embeddings of the joint modeling of both chromatin conformation and methylation of the sn-m3C-seq data (Lee et al., 2019).

Higashi has five main components. (1) We represent the scHi-C dataset as a hypergraph, where each cell and each genomic bin are represented as cell node and genomic bin node, respectively. Each non-zero entry in the single-cell contact map is modeled as a hyperedge connecting the corresponding cell and the two genomic loci of that particular chromatin interaction (Fig. 1a). Importantly, this novel formalism integrates embedding and data imputation for scHi-C. (2) We train a hypergraph neural network based on the constructed hypergraph (Fig. S1). (3) We extract the embedding vectors of cell nodes from the trained hypergraph neural network for downstream analysis. (4) We use the trained hypergraph neural network to impute single-cell Hi-C contact maps with the flexibility to incorporate the latent correlations between cells to enhance overall imputation, enabling more detailed and reliable characterization of 3D genome features. (5) With a number of new computational strategies, we reliably compare A/B compartment scores and TAD-like domain boundaries across individual cells to facilitate the analysis of cell-to-cell variability of these large-scale 3D genome features and its implication in gene transcription. In addition, we have developed a visualization tool to allow interactive navigation of the embedding vectors and the imputed contact maps from Higashi to facilitate discovery. The details are described in Methods.

### Higashi embeddings reflect cell types and cellular states

We sought to demonstrate that Higashi effectively captures the variability of 3D genome structures from the sparse scHi-C data with the embeddings. We first tested our method on three scHi-C datasets with multiple cell types or known cell state information at 1Mb resolution. These datasets include: 4DN sci-Hi-C dataset (Kim et al., 2020), Ramani et al. dataset (Ramani et al., 2017), and Nagano et al. dataset (Nagano et al., 2017). See Methods for data processing and Tables S1 and S2 for statistics of theses datasets. After training, the Higashi embeddings are projected to a two-dimensional space with UMAP (McInnes et al., 2018) for visualization. We found that the Higashi embeddings exhibit clear patterns that correspond to the underlying cell types and cellular states (Fig. S2a-c).

We then quantified the effectiveness of the embeddings by various evaluation settings and made direct comparisons to three existing scHi-C embedding methods, HiCRep/MDS (Liu et al., 2018), scHiCluster (Zhou et al., 2019), and LDA (Kim et al., 2020) (Supplementary Methods B.1). The quantitative results based on unsupervised evaluation suggest that the Higashi embeddings consistently outperform other methods (Fig. 1b). our evaluation shows that the Higashi embeddings can consistently achieve the best performance on scHi-C datasets with either categorical cell types or continuous cell states under various evaluation settings (Fig. S2d-f). Although all results in this section are based on the embedding with dimension size 64, our sensitivity analysis on the embedding dimension shows that Higashi is more robust to the choice of dimension size (Supplementary Results A.1 and Fig. S4a).

The emerging new technologies that jointly profile chromosome conformation and other epigenomic features have provided unique opportunities to directly analyze 3D genome structures and other modalities at single-cell resolution (Lee et al., 2019; Li et al., 2019). Higashi has the versatility to incorporate the co-assayed signals into the hypergraph representation learning framework as compared to separate analysis of two modalities, thereby taking full advantage of the co-assayed data (see Methods). We applied Higashi to a recently generated co-assayed dataset called single-nucleus methyl-3C (sn-m3C-seq) that jointly profiles Hi-C and DNA methylation in individual human prefrontal cortex cells (Lee et al., 2019). We found that the Higashi embeddings trained only on scHi-C (referred to as “Higashi (hic)”) can already resolve complex cell types in this dataset (Detailed results will be discussed in a later section). Importantly, when using Higashi to jointly model both signals (the embeddings referred to as “Higashi (joint)”), it reaches the overall best performance as compared to the embeddings based on only one modality (Fig. 1c). Higashi (joint) shows clearer patterns in the UMAP with cells being aggregated according to their cell types (Fig. 1d). Note that here the co-assayed methylation profiles are not part of the input but serve as the targets to approximate (see Methods).

Taken together, these results demonstrate that the Higashi embeddings effectively capture the cell-to-cell variability of 3D genome structures based on scHi-C data to reflect the underlying cellular states. In addition, the unique capability of Higashi for the joint modeling of both scHi-C and methylation profiles further enhances the scHi-C embeddings.

### Higashi robustly imputes scHi-C contact maps

In addition to dimension reduction of scHi-C data for cell type identification, Higashi can also impute sparse scHi-C contact maps. Here, we sought to demonstrate the imputation accuracy with several evaluations. For comparisons, we included the imputed results from scHiCluster. Note that scHiCluster represents each scHi-C contact map as an individual graph, whereas Higashi represents the whole scHi-C dataset as a hypergraph, allowing imputation to be potentially coordinated across different cells. Specifically, in Higashi, when imputing the contact map of cell *i*, its *k*-nearest neighbors in the embedding space would contribute to the imputation by taking advantage of their latent correlations (see Methods). To demonstrate the advantages of this important design employed in Higashi, we included the imputed results from Higashi with *k* as 0 and 4 (referred to as “Higashi(0)” and “Higashi(4)”, respectively). Importantly, we performed sensitivity analysis on the hyperparameter *k* and showed that Higashi is highly robust to the choice of *k* (Supplementary Methods A.1 and Fig. S4b).

We developed a novel simulation evaluation method to make use of the multiplexed 3D genome imaging data, which provides high-resolution physical views of 3D organization of genomic loci in individual cells (Bintu et al., 2018). Specifically, we turned the imaging data of a 2.5Mb region on chr21 from 11,631 cells at 30Kb resolution into scHi-C contact maps with various simulation coverage (Supplementary Methods B.2). We found that Higashi(0), i.e., no information sharing among different cells, can already consistently outperform scHiCluster. In addition, we found that Higashi(4) improves the imputation most significantly (30%-43% improvement on the median similarities across multiple metrics on the dataset with the lowest coverage). To illustrate why using neighboring cells in the embedding space improves imputation, we show a typical example from the simulated data with contact maps before and after imputation (Fig. 2 and Fig. S6). Consistent with the quantitative evaluation, Higashi(4) reveals the clearest patterns and especially domain boundaries across all coverage (Fig. 2 and Fig. S6). Importantly, the neighboring cells in the embedding space that contribute to the imputation indeed have similar 3D chromatin interactions compared with the selected cell, while the furthest cells do not. We carried out a similar set of evaluation using the more recent multiplexed imaging data of 3D genome structure (Su et al., 2020) (3,029 simulated contact maps of chr2 at 1Mb resolution) and reached the same conclusion of Higashi’s clear advantage (22%-50% improvement on the median similarities across multiple metrics on the dataset with the lowest coverage; see Fig. S7).

**Figure 2:**
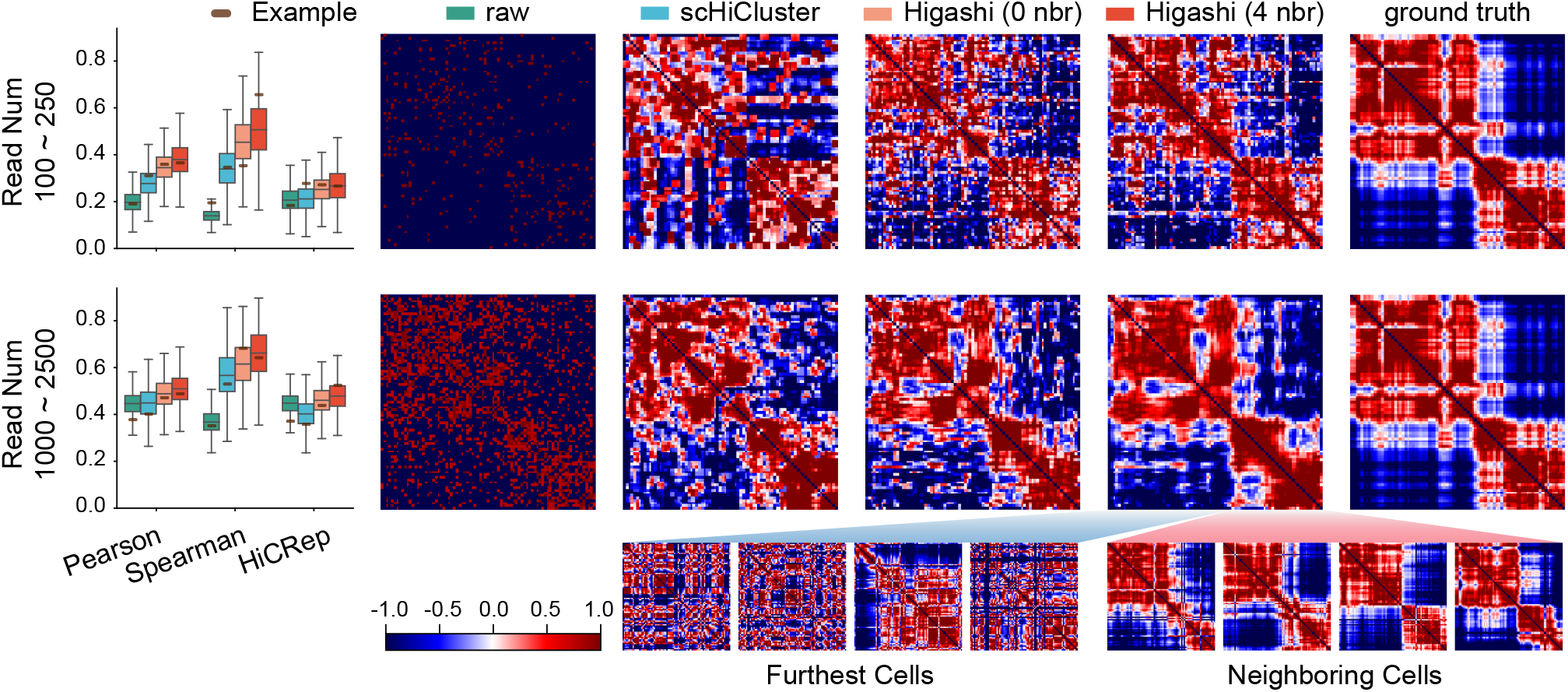
Evaluation and visualization of different imputation methods on scHi-C data simulated from multiplexed STORM 3D genome imaging data (Bintu et al., 2018). For Higashi, results by using information from four neighboring cells (4 nbr) or without using neighboring cell information (0 nbr) in the embedding space are both included. Each row corresponds to one set of simulation data with a chosen range of read numbers. The boxplots illustrate the quantitative evaluation of the similarities between the raw (input), the scHiCluster enhanced, and the Higashi enhanced contact maps versus the ground truth (inverse distance map). The heatmaps visualize the contact map before and after imputation as well as the ground truth. The contact maps of both the neighboring cells (in the embedding space) that contribute to the imputation and the cells that are furthest (in the embedding space) are shown. See also Fig. S6.

We performed additional evaluation via downsampling the existing scHi-C datasets with relatively higher coverage (Supplementary Methods B.2). We used the WTC-11 scHi-C dataset (personal communication with Bing Ren) of chr1 at 1Mb resolution and downsampled the sequencing reads of each cell at different rate (Methods; Table S1 and Table S4). We again observed clear advantage of Higashi for imputation with the strongest performance achieved by Higashi(4) (consistent advantages with up to 89% improvement on the distance stratified Spearman correlation; Fig. S8).

Together, these evaluations demonstrate that Higashi achieves much improved imputation of scHi-C contact maps robustly. The performance is further empowered by the unique mechanism of sharing information among neighboring cells in the embedding space. The improved imputation enables more reliable analysis of 3D genome structural features of each individual cell with higher accuracy.

### Higashi reveals compartmentalization variability at single-cell resolution

Next, we explored how the enhanced contact maps produced by Higashi facilitate new multiscale 3D genome analysis at single-cell resolution. A/B compartments reflect large-scale chromosome spatial segregation with distinct connections to genome function (Lieberman-Aiden et al., 2009). To date, little progress has been made for systematic A/B compartment annotation using scHi-C data primarily because of the data sparseness. Here, we applied Higashi to impute the WTC-11 scHi-C data at 50Kb resolution (see examples of the imputation results in Fig. S9). We designed a new method to calculate continuous compartment scores such that the scores are directly comparable across the cell population and reflect detailed cell-to-cell variation (Supplementary Methods B.3).

Fig. 3a shows the merged correlation matrices (Pearson correlation of the merged contact maps) before and after Higashi imputation as well as the compartment scores from the pooled scHi-C (i.e., bulk) and the single-cell compartment scores of chr21. After imputation, the merged scHi-C correlation matrix has much clearer checkerboard patterns that correspond to A/B compartments. The calculated single-cell compartment scores are overall consistent with the bulk compartment scores while showing cell-to-cell variability. Note that we identified one cluster of cells in the heatmap that have distinct patterns and are likely near the mitosis stage (marked with “^*^” in the bottom panel of Fig. 3a).

**Figure 3:**
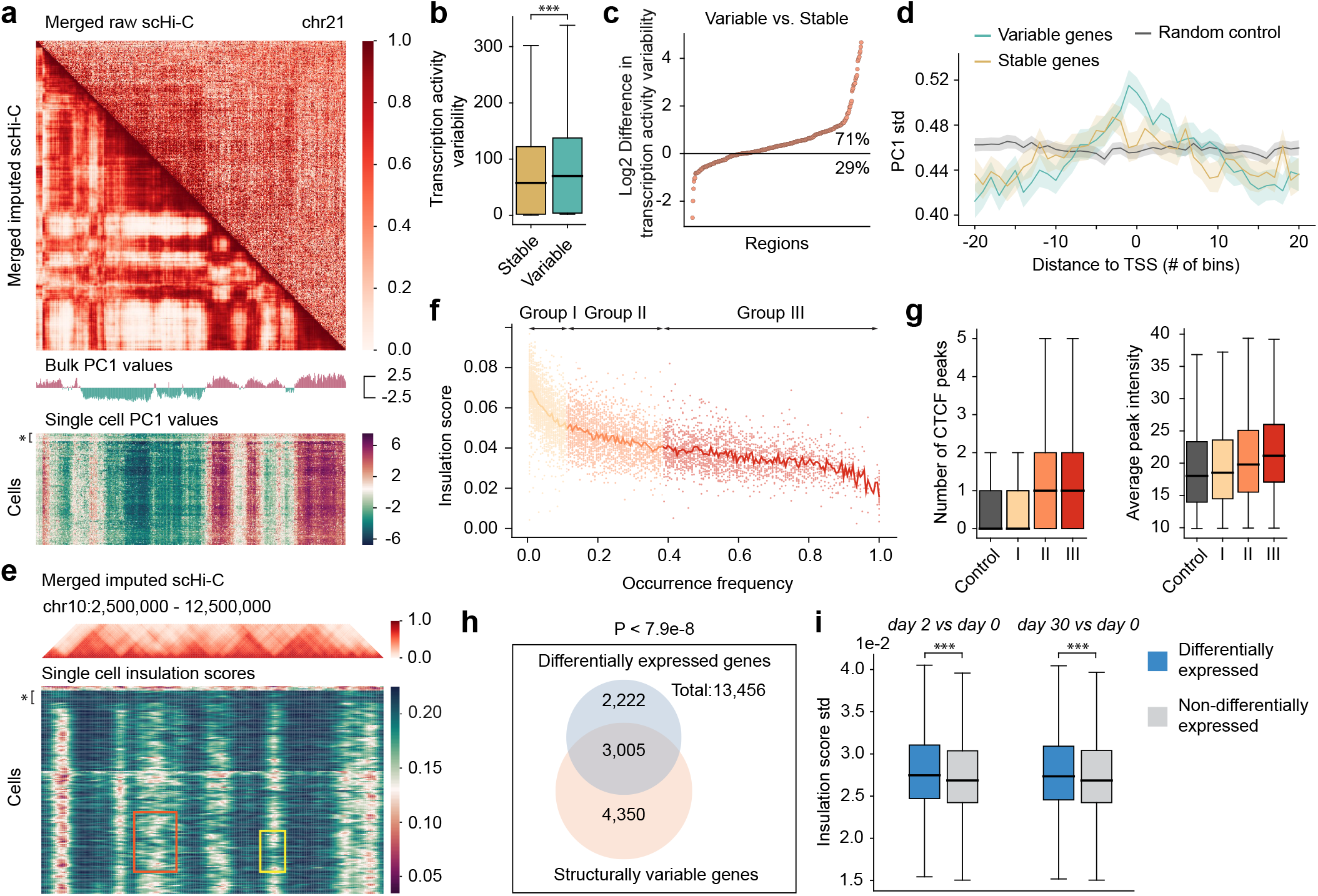
Higashi enables detailed characterization of 3D genome features and their connections to gene transcription at single-cell resolution. **a**. Compartment score annotations for WTC-11 scHi-C data at 50Kb resolution. The merged scHi-C correlation matrix on chr21 before and after imputation as well as compartment scores called from the pooled WTC-11 scHi-C data and each single-cell contact map are visualized. The cells that are likely near the mitosis stage are marked with “^*^” in the single-cell PC1 heatmap. **b**. Global comparisons of transcriptional variability on regions with variable and stable compartment annotations (^***^ indicates *P*-value*<*1e-3). There are 5,075 genes that have stable single-cell compartment scores with average transcription activity variability equal to 86.0. There are 5,071 genes that have dynamic single-cell compartment score with average transcription activity variability equal to 77.4. The one sided t-test *P*-value = 1.34 *×* 10^*−*7^. **c**. Log2 difference of transcriptional variability of genes with variable versus stable compartment annotations within a Mb scale window. **d**. Visualization of standard deviation of compartment scores around genes with variable or stable transcriptional level. In **b**,**c**,**d**, the transcriptional variability is quantified as the coefficient of variation of the imputed scRNA-seq data. **e**. TAD-like domain boundary calling for WTC-11 scHi-C at 50Kb resolution. The merged scHi-C contact maps at chr10:2,500,000-12,500,000 and the calculated insulation scores are shown. The cells that are likely near the mitosis stage are marked with “^*^” in the single-cell insulation score heatmap. Regions that represent the present/absent dynamics of single-cell domain boundaries are marked with a yellow box. Regions that represent the sliding dynamics of single-cell domain boundaries are marked with an orange box. **f**. Scatter plot of the single-cell insulation scores versus the occurrence frequency in the cell population of shared domain boundaries. For each cell, only the insulation scores of presented shared boundaries are visualized, i.e., each dot corresponds to a single-cell domain boundary. **g**. CTCF binding at domain boundaries from different occurrence frequency groups. **h**. Venn diagram of the overlap between genes near variable domain boundary in WTC-11 (light red) and differentially expressed genes during cell differentiation (light blue). **i**. Comparison of cell-to-cell variability of insulation scores between differentially expressed genes (DEGs) and non-DEGs. The high variance of insulation scores of DEGs indicates that the DEGs are enriched near domain boundaries with higher variability (^***^ indicates *P*-value*<*1e-3). Day 2 vs. day 0: 3,205 DEGs and 10,262 non-DEGs with mean insulation score standard deviation equal to 2.83 *×* 10^*−*2^ and 2.74 *×* 10^*−*2^, respectively. One-sided t-test *P*-value = 2.23 *×* 10^*−*9^. Day 30 vs. day 0: 4,308 DEGs and 9,159 non-DEGs with mean insulation score standard deviation equal to 2.80 *×* 10^*−*2^ and 2.74 *×* 10^*−*2^, respectively. One-sided t-test *P*-value = 4.16 *×* 10^*−*6^.

We explored the connection between the variability of compartment scores across the cell population and the transcriptional activity in different cells. We compared the compartment scores with the scRNA-seq from WTC-11 (Friedman et al., 2018). For this analysis, the cells that are likely near the mitosis stage were removed. For each gene, the transcriptional variability was calculated using the coefficient of variation (CV) (Supplementary Methods B.4). We quantified the compartment variability as the standard deviation of the single-cell compartment scores and further classified expressed genes as compartment variable or stable with a cutoff 50% based on the quantile. Compared with the transcriptional variability within these two groups (Fig. 3b), we observed that the genes in more variable compartments have higher transcriptional variability (*P*-value*<*0.001). We then used the 50Mb window resolution to assess if such structure-function variability correlation can also be observed at a finer scale. We used a 50Mb sliding window with a 1Mb step size on each chromosome and calculated the log difference of the median transcriptional variability between the variable and stable compartment regions within this window. As shown in Fig. 3c, among all windows, 71% of them follow the trend that genes in compartment variable regions have higher transcriptional variability. As a comparison, ∼76% of the genomic windows exhibit that the bulk compartment A correlates with higher expression levels (Lieberman-Aiden et al., 2009) (Fig. S11d). In addition, we made a step further to increase the resolution to individual genes. We classified genes as locally variable or stable by identifying the local minima/maxima of the transcriptional variability. We found that for the genes that are locally variable in terms of transcription, their compartment variability scores also tend to be the local maximum (Fig. 3d).

To confirm the robustness of these observations, in addition to using CV to measure transcriptional variability, we used another metric based a variance stabling algorithm (Supplementary Methods B.4) and reached similar observations (Fig. S11). These results further demonstrate the reliability of Higashi imputations, revealing cell-to-cell variability of compartment scores that are also functionally correlated.

### Higashi unveils TAD-like domain boundary variation across single cells

Recent work based on multiplexed STORM imaging of chromatin conformation revealed the existence and cell-to-cell variability of TAD-like structures in single cells (Bintu et al., 2018). However, the identification of TAD-like domains remains extremely challenging for sparse scHi-C data. We developed a new approach to reveal TAD-like domain boundary variability from single cells based on the Higashi imputations (Supplementary Methods B.5 and B.6). The analysis was conducted on the WTC-11 scHi-C dataset at 50Kb resolution.

We calculated single-cell insulation scores in which the local minima correspond to TAD-like domain boundaries (Crane et al., 2015) (Fig. 3e). We again observed a cluster of cells likely near the mitosis stage showing unidentifiable domain boundaries (marked with “^*^” in the bottom panel of Fig. 3e). We also observed that the local minima of the single-cell insulation scores often center around the domain boundaries observed in the merged imputed scHi-C, while the exact locations of the single-cell boundaries vary across the cell population (Fig. 3e). The dynamics of the single-cell domain boundaries have two main patterns: (1) present/absent across the population (marked with a yellow box in Fig. 3e); and (2) sliding along the genome (marked with an orange box in Fig. 3e). The first pattern reflects that a domain boundary does not occur in all cells. The second pattern manifests the shift of domain boundary along the genome, suggesting more gradual cell-to-cell variability. Comparison with scRNA-seq following the same approach used for single-cell compartment scores reached similar conclusions that domain boundary variability is strongly correlated with transcriptional variability at different scales (Fig. S11e-j).

Next, we made direct comparisons of TAD-like domain boundaries (Supplementary Methods B.6). As shown in Fig. 3f, where each dot corresponds to a single-cell domain boundary, we observed a negative correlation between the occurrence frequency of a domain boundary with its median single-cell insulation scores. This suggests that the more stable domain boundaries (i.e., higher occurrence frequency) from the cell population tend to be “stronger” boundaries in single cells associated with lower insulation scores. We also found positive correlation between the occurrence frequencies of domain boundaries and the number of CTCF binding peaks as well as the average CTCF peak intensity in the boundaries (Fig. 3g and Supplementary Results A.2). This result is consistent with the observation based on multiplexed STORM imaging (Bintu et al., 2018).

As an iPS cell type, WTC-11 can undergo cell differentiation. We identified differentially expressed genes (DEGs) from a scRNA-seq dataset of WTC-11 cells at 5 differentiation stages (Friedman et al., 2018) (Supplementary Methods B.7). Using hypergeometric test, we found that DEGs are over-represented in genes located near more variable domain boundaries in WTC-11 (top 50% of the insulation score std, *P*-value ≤ 7.9 × 10^*−*8^) (Fig. 3h). In addition, we compared the variability of insulation scores between DEGs and non-DEGs and found that DEGs have markedly higher standard deviation (one-sided t-test, *P*-value*<*0.001) (Fig. 3i). This suggests that the cell-to-cell variability of domain boundaries in WTC-11 may indicate important functional implication in cell differentiation.

Taken together, by analyzing the TAD-like domain boundaries across single cells enabled by Higashi, we revealed the correlation between domain boundary variability and gene regulation at single-cell resolution. These results further demonstrate the robustness and applicability of Higashi to facilitate the analysis of structure-function connections of 3D genome organization in single cells.

### Higashi reveals cell type-specific 3D genome features in human prefrontal cortex

To demonstrate Higashi’s ability to analyze single-cell 3D genome structures for complex tissues, we applied it to the aforementioned single-nucleus methyl-3C (sn-m3C-seq) data from the human prefrontal cortex (Lee et al., 2019). In this section, we present results from the Higashi framework trained only by the chromatin conformation information in sn-m3C-seq to evaluate its unique strength in analyzing scHi-C data.

We found that the Higashi embeddings (with scHi-C only) are able to resolve the differences among the neuron subtypes (separating Pvalb, Sst, Vip, Ndnf, L2-3, and L4-6), while maintaining clear separation with non-neuron cell types (Fig. 4a-b). This suggests that, empowered by Higashi, scHi-C alone has sufficient information to distinguish complex neuron subtypes. In contrast, scHiCluster cannot clearly distinguish these neuron subtypes using scHi-C (Fig. 5c in Lee et al. (2019)). Note that such advantage is confirmed by quantitative evaluation where the Higashi embeddings from scHi-C (“Higashi (hic)”) outperform the scHiCluster embeddings (Fig. 1c). In addition, a recent approach has been proposed to separate neuron subtypes on a dataset based on Dip-C with much higher coverage per cell (Tan et al., 2021). However, we found that for the sn-m3c-seq dataset, the method developed in Tan et al. (2021) cannot distinguish neuron subtypes (Fig. S13 and Supplementary Results A.3), further confirming the advantages of Higashi.

**Figure 4:**
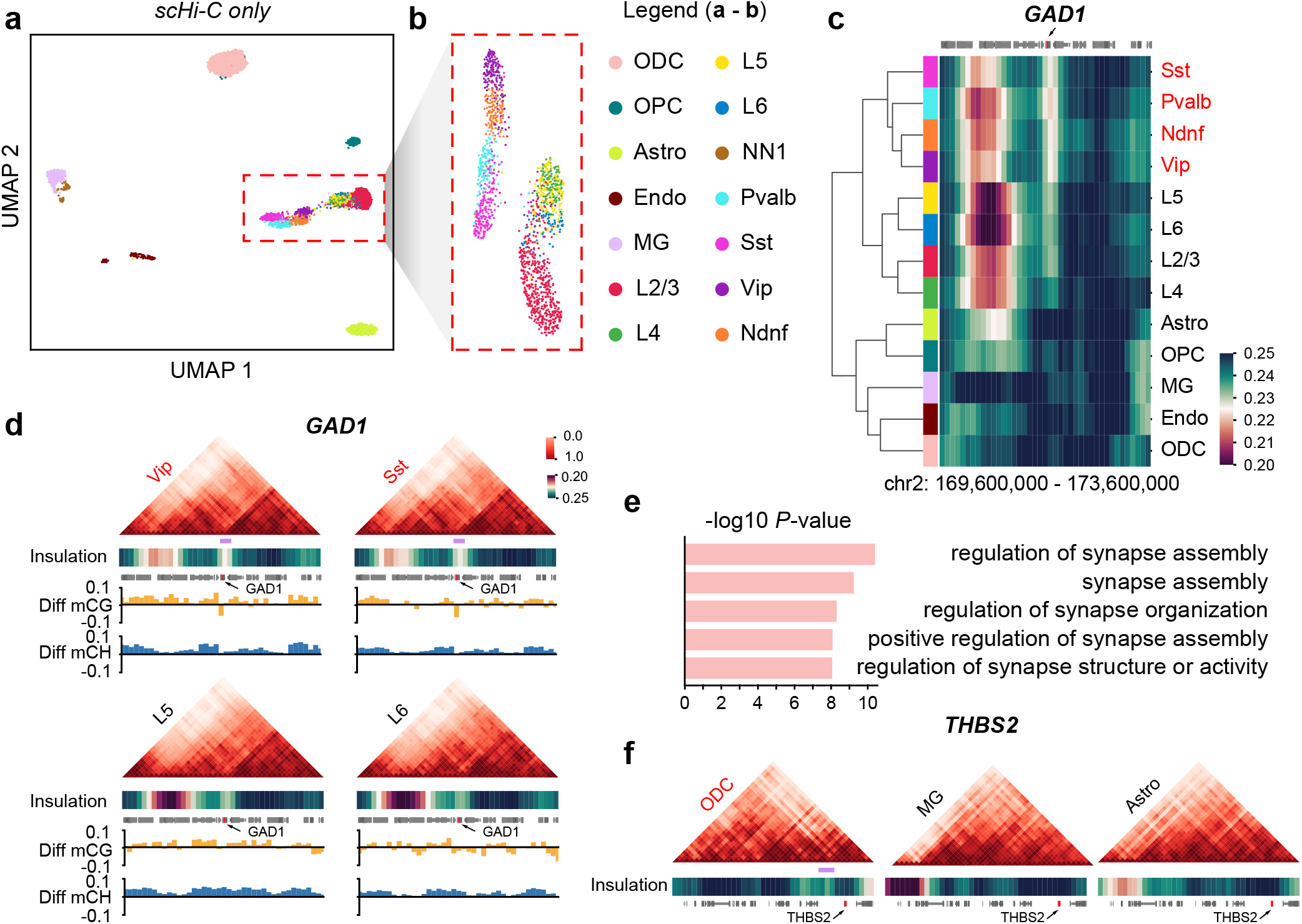
Higashi identifies complex cell types and cell type-specific TAD-like domain boundaries using scHi-C data from human prefrontal cortex. **a**. UMAP visualization of the Higashi embeddings using scHi-C only. **b**. UMAP visualization of the Higashi embeddings of the neuron subtypes in **(a). c**. Hierarchical clustering based on the average single-cell insulation scores of the flanking regions (+/-2Mbp) of the marker gene GAD1 for inhibitory neuron subtypes Sst, Pvalb, Ndnf, and Vip. Note that the single-cell insulation scores are calculated based on the Higashi imputed contact maps trained using only scHi-C data. **d**. Pooled imputed contact maps, average single-cell insulation scores, and methylation profiles of the same region in **(c)** for selected cell types. The methylation profile is calculated as the average CG/non-CG methylation percentage of a specific cell type minus the average CG/non-CG methylation percentage of the whole population. Light purple bar shows a TAD-like domain boundary specific to inhibitory neuron subtypes. **e**. Top five enriched gene ontology (GO) terms near ODC-specific TAD-like domain boundaries. **f**. Pooled imputed contact maps, insulation scores, and methylation profiles near the gene THBS2, which is in four of the top five most enriched GO terms with ODC-specific high expression. Light purple bar shows an ODC-specific TAD-like domain boundary. Cell type abbreviations in the legend (consistent with Lee et al. (2019)): L2/3, L4, L5 and L6: excitatory neuron subtypes; Ndnf, Vip, Sst, and Pvalb: inhibitory subtypes; Astro: astrocyte; ODC, oligodendrocyte; OPC, oligodendrocyte progenitor cell; MG, microglia; NN1, non-neuronal cell type 1; Endo, endothelial cell.

Next, we sought to identify cell type-specific TAD-like domain structures with the Higashi imputed contact maps. We found that the TAD-like domain boundaries show cell type-specific correlations with important known marker genes. For instance, the single-cell insulation scores of the region surrounding the transcription start site (TSS) of the maker gene GAD1 in inhibitory neuron subtypes reflect much stronger TAD-like domain boundaries (Fig. 4c). Note that such cell type-specific patterns are obscured in the pooled population contact maps (Fig. S14a). The cell type-specific domain boundary pattern is further manifested by comparison to the contact maps and methylation profiles (Fig. 4d and Fig. S15; light purple bars indicate cell type specific domain boundaries). In addition, we found that SULF1, which is a marker gene to distinguish subtypes L6 from the rest excitatory neuron subtypes (L2/3, L4, L5), has strong correlation with the surrounding cell type-specific TAD-like domain boundaries and methylation profiles (Fig. S14b, Fig. S16). Specifically, the TAD-like domain boundary is present in 93.2% of L6 cells but only in 65.3% of the rest excitatory neuron subtypes. These results provide new insights into the marker gene regulation of human prefrontal cortex cell types and the connection between 3D genome structure and function.

We next asked whether the genes near cell type-specific TAD-like domain boundaries identified by Higashi have distinct functional roles. We found that genes close to the ODC (oligodendrocyte) specific domain boundaries (784 in total) are strongly enriched with synapse-related GO terms as top hits (Fig. 4e; using GREAT (McLean et al., 2010)), suggesting the important role of ODC-specific domain boundaries in regulating synaptic functions (Allen and Lyons, 2018). To further analyze the connection between the ODC-specific domain boundaries and the regulation of the nearby genes, we further investigated the gene THBS2 which appears in four of the top five GO term categories we identified. THBS2 is known to be expressed in glial cells and is key to the regulation of synaptic functions (Allen and Eroglu, 2017). The visualization of the pooled contact maps of the 4Mb region surrounding THBS2 shows that ODCs have a TAD-like domain boundary upstream of the TSS of THBS2 (light purple bar in Fig. 4f, Fig. S17), which can be elucidated by single-cell insulation scores of this region (Fig. S14c). Importantly, the TAD-like domain boundary near THBS2 is obscured in the insulation score calculated from the population contact map (Fig. S14c), further supporting the importance of scHi-C analysis enabled by Higashi. Note that THBS2 has cell type-specific high expression in ODC (fold-change 8.6 compared with the population average) (Hawrylycz et al., 2012). Therefore, the ODC-specific TAD-like domain boundaries may offer new perspectives for understanding the cell type-specific gene regulation of THBS2.

Taken together, these results demonstrate the distinct ability and advantages of Higashi to effectively identify cell types and cell type-specific 3D genome features in complex tissues using scHi-C data. Crucially, this analysis shows the strong potential of Higashi in revealing cell type-specific TAD-like domain boundaries, greatly facilitating the analysis of the roles of 3D genome structure in regulating cell type-specific gene function.

## Discussion

In this work, we developed Higashi for multiscale and integrative scHi-C analysis. Our extensive evaluation demonstrated the advantage of Higashi over existing methods for both embedding and imputation. Additionally, enabled by the improved data enhancement of scHi-C contact maps, we developed new methods in Higashi to systematically analyze variable multiscale 3D genome features (A/B compartment scores and TAD-like domain boundaries), revealing their important implications in gene transcription. By applying to a scHi-C dataset from human prefrontal cortex, Higashi is able to identify complex cell types and reveal cell type-specific TAD-like boundaries that have strong connections to cell type-specific gene regulation.

The key algorithmic innovation of Higashi is the transformation of scHi-C data into a hypergraph, which has unique advantages as compared to existing methods. First, this transformation preserves the single-cell precision and 3D genome features from scHi-C. Second, modeling the whole scHi-C datasets as a hypergraph instead of modeling each contact map as individual graphs allows information to be coordinated across cells to improve both embedding and imputation by taking advantage of the latent correlations between cells. Third, although we mainly focused on scHi-C data, the hypergraph representation in Higashi is highly generalizable to other single-cell data types. As a proof-of-principle, we showed that Higashi can be extended to analyze co-assayed scHi-C data with methylation in an integrated manner, showing markedly improved performance as compared to separate analysis of the two modalities. In addition, to achieve more comprehensive delineation of 3D genome organization at single-cell resolution, Higashi can be potentially extended to analyze single-cell assays of higher-order chromatin structures, e.g., the recently developed scSPRITE (Arrastia et al., 2020) that probes multi-way chromatin interactions.

It is anticipated that many exciting new scHi-C data will become widely available in the coming years. Higashi is a robust and powerful new framework that can serve as an essential tool to effectively analyze such data, providing uniquely new insights into nuclear genome structure and function at single-cell resolution in various biological contexts.

## Methods

### scHi-C data and other genomic data processing

In this work, we used several publicly available single-cell Hi-C datasets. We refer to them as: Ramani et al. (Ramani et al., 2017), Nagano et al. (Nagano et al., 2017), 4DN sci-Hi-C (Kim et al., 2020) (4DN Data Portal: 4DNES4D5MWEZ, 4DNESUE2NSGS, 4DNESIKGI39T, 4DNES1BK1RMQ, 4DNESTVIP977). We also used a new scHi-C dataset generated from the WTC-11 iPSC line (4DN Data Portal: 4DNESF829JOW, 4DNESJQ4RXY5).

For all scHi-C datasets, we only kept the cells with more than 2,000 read pairs that have genomic span greater than 500Kb. At a given resolution, we define the number of contacts per cell as the number of interaction pairs (read count) assigned to the non-diagonal entries of the intrachromosomal contact maps. The Ramani et al. dataset and the 4DN sci-Hi-C dataset used single-cell combinatorial indexed Hi-C (sci-Hi-C). After filtering, the Ramani et al. dataset contains 620 cells of four human cell types (GM12878, HAP1, HeLa, K562) with 7,800 median contacts per cell, while the 4DN sci-Hi-C dataset contains 6,388 cells of five human cell types (GM12878, H1ESC, HAP1, HFFc6, IMR90) with 3,800 median contacts per cell. The Nagano et al. dataset used a different protocol with 1,171 cells and 56,800 median contacts per cell. The WTC-11 scHi-C data (188 cells in total) was generated using single-nucleus Hi-C with 144,800 median contacts per cell. The co-assayed single-cell methylation and Hi-C dataset (sn-m3C-seq) was from Lee et al. (2019). We followed the same processing pipeline as sn-m3C-seq for processing the methylation signals.

We also used other publicly accessible genomic datasets in this work. The scRNA-seq of WTC-11 was from Friedman et al. (2018). The details on calculating transcriptional variability based on scRNA-seq can be found in Supplementary Methods B.4. We also analyzed the CTCF binding near the identified single-cell TAD-like domain boundaries in WTC-11 cells. We used the WTC-11 CTCF ChIA-PET data (4DN Data portal: 4DNES8MZ76GP) and called peaks based on the singleton reads from the dataset following the ENCODE ChIP-seq peak calling pipeline (Moore et al., 2020). Specifically, peaks were generated for individual replicates and merged by keeping only the reproducible peaks.

### Overall structure of the hypergraph neural network in Higashi

A *hypergraph G* is a generalization of graph and can be formally defined as a pair of sets *G* = (*V, E*), where *V* = {*v*_*i*_} represents the set of nodes in the graph, and 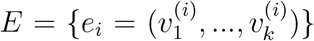 represents the set of hyperedges. For any hyperedge *e* ∈ *E*, it connects two or more nodes (|*e*| ≥ 2). Both nodes or hyperedges can have attributes reflecting the associated properties such as node type or the strength of a hyperedge. The *hyperedge prediction problem* aims to learn a function *f* that can predict the probability of a group of nodes (*v*_1_, *v*_2_, …, *v*_*k*_) forming a hyperedge or the attributes associated with the hyperedge. For simplicity, we refer to both cases as predicting the probabilities of forming a hyperedge.

The core part of Higashi is a hypergraph representation learning framework, extending our recently developed Hyper-SAGNN (Zhang et al., 2020), that models higher-order interaction patterns from the hypergraph constructed from the scHi-C data. The overall structure of the hypergraph neural network is illustrated in Fig. S1. We use *x*_*i*_ to represent the attributes of node *v*_*i*_. The input to the model is a triplet, i.e., (*x*_1_, *x*_2_, *x*_3_) consisting of attributes of one cell node and two genomic bin nodes. We do not differentiate between these two types of nodes in this section for simplicity. Each node within a triplet passes through a neural network (NN), respectively, to produce (*s*_1_, *s*_2_, *s*_3_), where *s*_*i*_ = NN_1_(*x*_*i*_). The structure of NN_1_ used in this work is a position-wise feed-forward neural network with one fully connected layer. By definition, each *s*_*i*_ remains the same for node *v*_*i*_ independent to the given triplet, and is thus called the “static embedding”, reflecting the general topological properties of a node in the given hypergraph. In addition, the triplet as a whole also passes through another transformation, leading to a new set of vectors (*d*_1_, *d*_2_, *d*_3_), where *d*_*i*_ = NN_2_(*x*_*i*_ | (*x*_1_, *x*_2_, *x*_3_)). The structure of NN_2_ will be discussed later. The definition of *d*_*i*_ depends on all the node features within this triplet that reflects the specific properties of a node *v*_*i*_ in a particular hyperedge and is thus called the “dynamic embedding”.

Next, the model utilizes the difference between the static and dynamic embeddings to produce ŷ_*i*_ by passing the Hadamard power of *d*_*i*_ − *s*_*i*_ to a fully-connected layer. Additional features, including the genomic distance between the bin pair, one hot encoded chromosome ID, and also the total read number per cell, are concatenated and sent to a multi-layer perceptron with output ŷ_ext_. All the output ŷ_*i*_ and ŷ_ext_ are further aggregated to produce the final result ŷ, i.e., the predicted probability for this triplet to be a hyperedge:

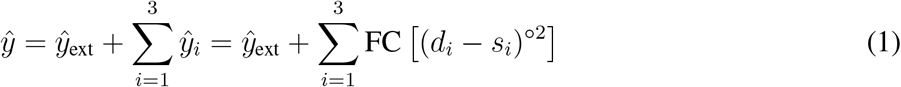

where FC is the fully-connected layer.

In the following sections, we describe how the node attributes are generated, the structure of NN_2_, the model training, and how we incorporate co-assayed signals into Higashi.

### Node attribute generation in Higashi

As mentioned, the input to the hypergraph neural network model is a triplet consisting of attributes of one cell node and two genomic bin nodes. For the bin nodes, we use the corresponding rows of the merged scHi-C contact maps as the attributes. For the cell nodes, we calculate a feature vector based on its single-cell contact maps as its attributes. This process is as follows:

1. Each contact map is normalized based on the total read count.
2. Contact maps are flattened into 1D vectors and concatenated across the cell population.
3. (optional SVD (singular value decomposition) is used to reduce dimensions for computational efficiency.
4. The corresponding row in the feature matrix is used as the attributes for the corresponding cell.

### Cell dependent graph neural network with self-attention for dynamic embeddings

Here, we introduce the neural network NN_2_ (mentioned above) that transforms the attributes of a node given a node triplet to the corresponding dynamic embeddings. In the original Hyper-SAGNN, this was accomplished by the multi-head self-attention layer (Vaswani et al., 2017) (see Zhang et al. (2020)). However, the representation capacity of using self-attention layers to calculate dynamic embeddings is constrained by the embedding dimensions and the depth of self-attention layers, which would lead to high computational cost and increased training difficulty.

To increase the expressiveness of this neural network for generating dynamic embeddings while maintaining small embedding dimensions and fewer layers, we developed a cell dependent graph neural network (GNN) (Hamilton et al., 2017) that transforms the attributes of bin nodes before passing to the self-attention layer. For a node triplet (*c*_*i*_, *b*_*j*_, *b*_*k*_), where *c*_*i*_ corresponds to a cell node and *b*_*j*_, *b*_*k*_ are bin nodes, a graph *G*(*c*_*i*_) (where both *b*_*j*_, *b*_*k*_ are nodes in it) is constructed by taking *c*_*i*_ as input. Details on the construction of *G*(*c*_*i*_), which is shared for all triplets that contain the cell node *c*_*i*_, is discussed in the next section. For each layer in the GNN, to generate the output vector for bin node *b*_*j*_, the information of its neighbors in the graph 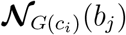 is aggregated:

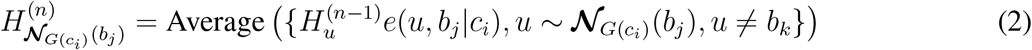

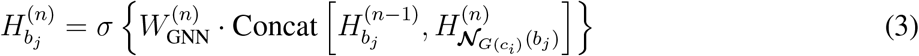

where 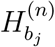 is the output vector of the node *v*_*i*_ at the *n*-th layer of the GNN and 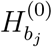 is the attributes of the node *b*_*j*_ before passing to the GNN. 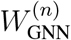 represents the weight matrix to be optimized at the *n*-th layer and *σ* is the non-linear activation function. We call this GNN cell dependent because the structure of the graph depends on the cell, although the weight matrix 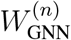 is shared across all cells. This cell dependent GNN can improve the expressiveness of the neural network by incorporating a large amount of single-cell information (contact maps) into the structure of the model instead of entirely relying on the embeddings of the cell nodes. The GNN is trained to reconstruct the interaction between a pair of bin nodes by using only information of themselves and their neighborhood (but not including each other). The attributes of both *b*_*j*_ and *b*_*k*_ are transformed by this cell dependent GNN into 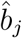 and 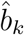, respectively, and the triplet of 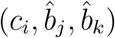 passes through the aforementioned self-attention layer to generate the final dynamic embeddings.

### Information sharing among cells with similar genome structures

Higashi has a unique capability for cells to share information with each other in the embedding space to enhance imputation by taking advantage of the latent correlations between cells. Specifically, we first train Higashi until convergence without the cell dependent GNN to allow the self-attention layer to capture cell-type specific information and reflect in the embeddings through back-propagation. We then calculate the pairwise distances of cell embeddings that indicate the similarities among cells. Given a hyperparameter *k*, we construct a graph *G*(*c*_*i*_) based on the contact maps of *c*_*i*_ and its *k*-nearest neighbors in the embedding space. It is crucial to clarify that, when we mention the neighbor of a cell 𝒩 (*c*_*i*_), we are referring to other cells that have small pairwise distances of embedding vectors instead of other nodes that have connections to the cell in the hypergraph. We name the contact maps of *c*_*i*_ as *M* (*c*_*i*_). The new *G*(*c*_*i*_) is constructed as the weighted sum of *M* (*u*), *u* ∈ {*c*_*i*_} ∪ 𝒩 (*c*_*i*_), where the weight is calculated based on the pairwise distance *d*(*u, c*_*i*_) in the embedding space, i.e.,

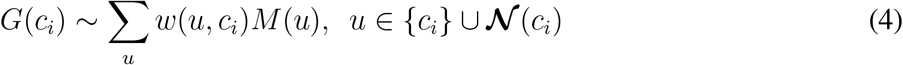

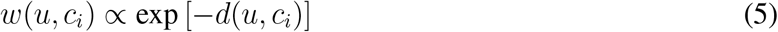

Each embedding is normalized by the maximum 𝓁^2^ norm. Note that although contact maps of different cells are mixed in this step, we do not mix the prediction results from different cells or directly use the mixed contact maps as output. This differentiates our method from the *k*-NN based smoothing methods fundamentally. The Higashi model is trained with only the observed interactions in each single cell, together with the interactions in cells that share overall similar structures serving as auxiliary information for synergistic prediction in a cell population.

### Model training details in Higashi

The hypergraph neural network within Higashi would produce a score ŷ for any triplet (*c*_*i*_, *b*_*j*_, *b*_*k*_). The neural network is trained to minimize the difference between the predicted score ŷ and the ground truth score *y*, indicating the probability of the pairwise interaction between bin nodes *b*_*j*_ and *b*_*k*_ in cell *c*_*i*_. In Higashi, we offer two choices of loss function for scHi-C datasets with different coverage. For scHi-C datasets with relatively low sequencing depths or the analysis resolution is high (hence fewer reads in each genomic bin), the model is trained with a binary classification loss (cross entropy) where the triplets corresponding to all non-zero entries in the single-cell contact maps are treated as positive samples and the rest are considered as the negative samples (i.e., *y*(*c*_*i*_, *b*_*j*_, *b*_*k*_) ∈ {0, 1}). The classification loss is:

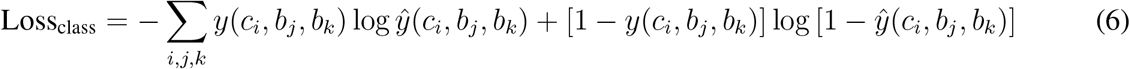

For datasets with relatively high sequencing depths or when the analysis resolution is low (hence more reads in each genomic bin), we further differentiate among the non-zero values by training the model with a ranking loss, which maintains consistent ranking of predicted scores versus the continuous ground truth scores (i.e., *y*(*c*_*i*_, *b*_*j*_, *b*_*k*_) ∈ ℝ). The ranking loss can be described as a binary classification problem aiming to identify the triplet with larger ground truth score in a pair of selected triplets. For simplicity, we denote two triplets as *t*_*i*_, *t*_*j*_ and the corresponding ground truth scores as *y*(*t*_*i*_), *y*(*t*_*j*_). The ranking loss is:

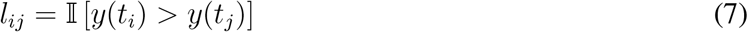

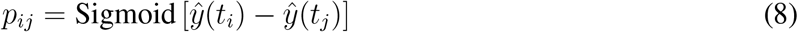

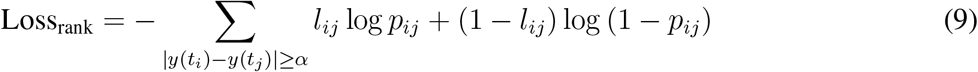

where *α* defines whether the order of *y*(*t*_*i*_), *y*(*t*_*j*_) can be reliably called and is set to 2 in this work. Note that *l*_*ij*_, *p*_*ij*_ are intermediate variables only used in this definition.

Using either the classification loss or the ranking loss requires negative samples in the training data. We designed an effective negative sampling approach. Specifically, at each epoch, we randomly sample a batch of triplets and make sure that these triplets do not overlap with the positive samples. To reflect the similarity of 3D genome structures of flanking genomic bins, we also exclude triplet (*c*_*i*_, *b*_*j*_, *b*_*k*_) from the negative samples if triplets such as (*c*_*i*_, *b*_*j*_ + 1, *b*_*k*_) belong to the positive samples. The number of negative samples generated for each batch is guided by the sparsity of the input data. When studying a scHi-C data where *s*% of the contact map entries are zeros, for a batch of *n* positive triplets, min [*s/*(100 − *s*), 5] *n* negative samples will be generated. The number of negative samples is no more than 5 times of the number of positive samples for computational efficiency. The model is optimized by the Adam algorithm (Kingma and Ba, 2014) with learning rate of 1e-3. We train Higashi for 80 epochs with batch size of 192 or until the model converges on an individual validation set before 80 epochs.

### Incorporating co-assayed signals in Higashi

The unique design of Higashi allows joint modeling of co-assayed scHi-C and the corresponding one dimensional signals (e.g., from sn-m3C-seq (Lee et al., 2019)). We add an auxiliary task for Higashi by using the learned embeddings for cell nodes *c*_*i*_ to accurately reconstruct the co-assayed signals *m*_*i*_ through a multi-layer perceptron (MLP). The auxiliary loss term is added to the main loss function and optimized jointly. The model thus builds an integrated connection between chromatin conformation and the co-assayed signals, guiding the embedding of the scHi-C data, i.e.,

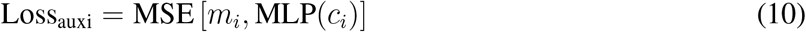

where MSE refers to the mean squared error between the ground truth of co-assayed signals and the estimate.

### Methods for the analysis of variable compartment and domain boundary from scHi-C

In Higashi, we have developed new strategies for reliable analysis of 3D genome features in different scales across the cell population. We developed a method to calculate continuous compartment scores for the imputed single-cell contact maps such that these scores are directly comparable across different cells in the population to assess variability (see Supplementary Methods B.3). For single-cell TAD-like domain boundary analysis, we developed a new calibration method with a novel optimization scheme to achieve comparative analysis of domain boundary variability from single cells based on insulation scores (see Supplementary Methods B.5 and B.6). These algorithms greatly enhance the analysis of variable multiscale 3D genome structures at single-cell resolution.

### Visualization tool for integrative scHi-C analysis

In Higashi, we developed a new visualization tool that allows interactive navigation of the scHi-C analysis results. Our tool enables the navigation of the embedding vectors and the imputed contact maps from Higashi in a user-friendly interface. Users can to select individual or group of cells of interest in the embedding space and explore the corresponding single-cell or pooled contact maps of interest. Fig. S18 shows a screenshot of the visualization tool. See the GitHub repository of Higashi for detailed documentation of this visualization tool: https://github.com/ma-compbio/Higashi.

### Code Availability

The source code of Higashi can be accessed at: https://github.com/ma-compbio/Higashi.

## Acknowledgements

The authors would like to thank Bing Ren for sharing the WTC-11 single-cell Hi-C data before publication, Yijun Ruan for making the WTC-11 CTCF ChIA-PET data available, Jingtian Zhou for suggestions on the sn-m3c-seq data analysis, and Yang Zhang for suggestions that improved the manuscript. The authors are also grateful to Bing Ren, Job Dekker, William Noble, Zhijun Duan, and other members of the NOFIC-AICS Collaborative Project Working Group and the Joint Analysis Working Group of the NIH 4DN Consortium for discussions and feedback. This work was supported in part by the National Institutes of Health Common Fund 4D Nucleome Program grants U54DK107965 (J.M.) and UM1HG011593 (J.M.) and National Institutes of Health grant R01HG007352 (J.M.). J.M. is additionally supported by a Guggenheim Fellowship from the John Simon Guggenheim Memorial Foundation.

## Author Contributions

Conceptualization, R.Z. and J.M.; Methodology, R.Z. and J.M.; Software, R.Z.; Investigation, R.Z., T.Z., and J.M.; Writing – Original Draft, R.Z. and J.M.; Writing – Review & Editing, R.Z., T.Z., and J.M.; Funding Acquisition, J.M.

