## Supplemental Information for "Multiscale and integrative single-cell Hi-C analysis with Higashi"

#### A Supplementary Results

##### A.1 Sensitivity analysis of hyperparameters in Higashi

Here, we discuss how we evaluated the robustness of Higashi to the hyperparameters in the model. We first tested Higashi's embeddings with regard to the embedding dimensions based on the Ramani et al. dataset. We trained Higashi as well as the baseline methods, scHiCluster (Zhou et al., 2019), HiCRep/MDS (Liu et al., 2018), and LDA (Kim et al., 2020), to produce embeddings with different dimensions. The details for the evaluation procedure can be found in Supplementary Methods B.1. As shown in Fig. S4a, among all four methods, Higashi achieves the highest scores and is the most robust to the embedding dimensions. Under the supervised setting, the embedding vectors produced by scHiCluster, HiCRep/MDS and LDA have a strong overfitting problem where the logistic regression classifier trained on 10% of the embeddings do not generalize well to the rest of the cells. However, the embeddings of Higashi do not encounter this problem with stable Micro-F1 and Macro-F1 scores across different embedding dimensions. Under the unsupervised setting, Higashi performs consistently well across all embedding dimensions. The performance quantified by ARI becomes even better for higher dimensional embeddings. The other three baseline methods achieve relatively stable ARI scores across different embedding dimensions as compared to their Micro-F1 and Macro-F1 scores in the supervised setting, but they either perform worse in general or are overall not as robust as Higashi. Notably, we observed that the method LDA can not achieve reasonable clustering without the utilization of UMAP (marked with LDA (UMAP) in Fig. S4a).

The second hyperparameter that we evaluated in Higashi is the number of nearest neighbors in the embedding space that coordinate the data imputation for a specific cell. As mentioned, one unique novelty of Higashi with the hypergraph formulation is the ability to allow information to be shared across different cells, which is controlled by the hyperparameter $k$. This parameter can be chosen based on the sparsity of the contact map to be imputed and also the number of cells within the dataset. For instance, if the coverage of the dataset is very limited, it is appropriate to have a larger $k$ to enhance the imputation. Moreover, if the number of cells within the dataset is large, a higher $k$ would also be desired because there is great probability that a cell in such a population shares similar 3D genome structure with other cells. In order to evaluate if Higashi is robust to the choice of $k$, we repeated the simulation evaluation based on the Bintu et al. (2018) data with $k = 0, 2, 4, 6$ and $\infty$ (using all other cells within the experiment), respectively. As shown in Fig. S4b, across different sampled read numbers, Higashi(2), Higashi(4), and Higashi(6) always achieve higher similarity scores with the ground truth compared with Higashi(0) and the baseline models. The difference in the performance among Higashi(2), Higashi(4), and Higashi(6) is minimal, strongly suggesting the robustness of Higashi to the hyperparameter $k$. We also included the extreme case where there is no limit on the number of neighboring cells involved in the imputation. We found that this will lead to a slight decrease in imputation accuracy (Fig. S4b). However, as compared to Higashi(0) and other baselines, Higashi($\infty$) still achieves higher or comparable performance.

We also repeated this evaluation on the simulated data based on the imaging data of the whole chr2 from Su et al. (2020). Similar to what we observed on the simulation data based on Bintu et al. (2018), the advantages of Higashi(2), Higashi(4), and Higashi(6) over other methods are significant with very little variance among them. Higashi($\infty$) decreases the imputation accuracy under certain evaluation metrics but still performs reasonably well as compared to the baseline models (Fig. S4c).

Taken together, we demonstrated that the unique design in Higashi to allow information to be shared between neighboring cells in the embedding space consistently improves data imputation with robust performance. The choice of hyperparameter $k$ affects the imputation results only slightly in Higashi with consistent advantages over other methods, further suggesting the robustness of the method.

##### A.2 Analysis of CTCF binding near single-cell TAD-like domain boundaries

From earlier work using bulk Hi-C, it is known that TAD boundaries are typically enriched with CTCF binding (Dixon et al., 2012). We sought to reveal the connection between CTCF binding and TAD-like domain boundaries from scHi-C. Based on the occurrence frequencies of the shared TAD-like domain boundaries in the cell population, we classified the shared boundaries into three groups by quantile: (1) group I corresponds to the dynamic domain boundaries; (2) group II corresponds to the moderately dynamic domain boundaries; and (3) group III corresponds to the relatively stable ones. We analyzed the CTCF binding within these different domain boundary groups. We used the peaks called from the CTCF ChIA-PET data in WTC-11 (see Methods). We found that both the number of CTCF binding peaks and the average peak intensity are positively correlated with the domain boundary occurrence frequencies, which is consistent with the observation in Bintu et al. (2018) based on multiplexed STORM imaging data (see Fig. 2g in main text). Specifically, the group I boundaries have less overlap with the CTCF binding. On the other hand, group II and III boundaries are more enriched with CTCF binding. Notably, group II boundaries, which appear in less than 50% of the cell population, still have similar CTCF binding compared to more stable ones. Instead of analyzing only the reproducible peaks, we repeated the analysis on each replicate and observed the same patterns (Fig. S12).

##### A.3 Analysis of using single-cell A/B compartment scores as embeddings

Tan et al. (2021) reported that after PCA the single cell A/B compartment scores (referred to as scA/B compartment scores) can be used as embeddings to identify cell types and subtypes on a Dip-C dataset with higher coverage. In the main text, we showed that by using only the chromatin conformation information of the sn-m3c-seq dataset as input, Higashi embeddings can already separate major neuron subtypes (Fig. 4a), providing major conceptual advance compared with previous work (Note that Lee et al. (2019) reported that chromatin conformation does not have enough information to identify complex neuron subtypes). In this section, we sought to test if scA/B compartment scores can be used as embeddings on the sn-m3c-seq dataset as well. It is worth noting that the sn-m3c-seq dataset from Lee et al. (2019) has lower coverage as compare to the Dip-C dataset from Tan et al. (2021).

In Tan et al. (2021), scA/B compartment scores are approximated by the CpG frequency. The scA/B compartment scores are further rank-normalized within each cell. Besides rank-normalization, we also tested another normalization method that is less sensitive and more robust, which is the quantile transform with the maximum number of quantiles chosen as 20% of the number of bins per cell. This would be equivalent to merging similar ranks into the same value in the rank-normalized scA/B compartment scores. The scA/B compartment scores calculated based on CpG frequency and normalized by these two approaches are referred to as "scA/B (cpg) - RankTie" and "scA/B (cpg) - Quantile", respectively. These scA/B compartment scores are further transformed through PCA with top 20 PCs kept for UMAP visualization (the same as Tan et al. (2021)). For comparisons, we also included the scA/B compartment scores calculated using the approach proposed in this work (see Supplementary Methods B.3 for details), which are further normalized with quantile transform (referred to as "scA/B (Higashi) - Quantile"). All scA/B compartment values are in 1Mb resolution.

As shown in Fig. S13, all embeddings based on the scA/B compartment scores cannot separate neuron subtypes as well as the Higashi embeddings (Fig. 4a), highlighting the advantages of Higashi for generating scHi-C embeddings. We also observed that the scA/B compartment scores calculated based on CpG frequency show apparent batch effects, whereas such batch effects are mitigated with scA/B (Higashi) (Fig. S13). This strongly suggests Higashi's ability in removing batch effects when imputing the contact maps. We also found better separation of inhibitory and excitatory neuron subtypes and overall better separation among non-neurons with the UMAP visualization of scA/B (Higashi) (Fig. S13).

Taken together, we demonstrate the clear advantage of Higashi embeddings in separating complex cell types. Additionally, our method for calculating single-cell compartment scores based on the Higashi imputed contact maps is more robust to the batch effects and can reflect the cell types more accurately.

### B Supplementary Methods

#### B.1 Evaluation procedure of the embedding vectors

In this work, we quantified the effectiveness of the embeddings from different methods (Higashi, HiCRep/MDS (Liu et al., 2018), scHiCluster (Zhou et al., 2019) and LDA (Kim et al., 2020)) with various supervised and unsupervised evaluation tasks. We kept the embedding dimensions and the resolution of the input scHi-C contact maps the same for all methods to make fair comparisons. The rest hyperparameters were kept as the default or the ones that were used in the original study for a specific dataset. For instance, for LDA, we set the maximum distance between a locus pair as 10Mb for the Ramani et al. and 4DN-sci-Hi-C datasets, and 25Mb for the Nagano et al. dataset, which was the same parameter setting in Kim et al. (2020). For sn-m3c-seq dataset (Lee et al., 2019), we also included the embeddings generated from the CG methylation (mCG) profile using Scanorama (Hie et al., 2019) for comparisons. The embedding dimension is kept the same as the other embeddings based on scHi-C for direct comparisons. Note that the dimension of mCG embeddings is different from what is used in Lee et al. (2019) to annotate the cell types. For the supervised setting, we trained a logistic regression classifier with 10% of the cells as training data and the remaining 90% cells as testing. The supervised setting is designed to mimic the common practice of confidently annotating a small fraction of cells with high coverage based on prior knowledge (e.g., marker genes). Because the value range of embeddings produced by different methods vary, which would affect the convergence as well as the numerical stability, we applied Z-score normalization before training the logistic regression classifier. Note that linear transformation on the embeddings would not change the solution of a linear classifier such as logistic regression. It only affects the convergence rate and the numerical stability of the algorithm for solving the optimal weights of the classifier. The performance for this multi-class classification task was measured by Micro-F1 and Macro-F1 scores. For the unsupervised setting, we applied $K$-means clustering on datasets with discrete cell states (such as the 4DN sci-Hi-C and Ramani et al. datasets used in this work) and calculated the Adjusted Rand Index (ARI) of the predicted labels against the cell types. Because Z-score normalization would change the weight of each dimension when calculating euclidean distances in $K$-means clustering, we did not apply Z-score normalization to each dimension separately but rather scaled the embeddings across different dimensions altogether with Z-score. For dataset with continuous cell states (the Nagano et al. dataset used in this work that represent states of continuous cell cycle), we used a metric called ACROC (Average Circular ROC) proposed in Liu et al. (2018) to evaluate the consistency of embedding vectors with circulatory states. All metrics used in this evaluation range from 0 to 1 with larger number indicating better performance. For comparison, all methods in our evaluation generated 64-dimensional embeddings for all evaluation metrics except ACROC since it is defined in the two-dimensional space. When calculating the ACROC score, both scHiCluster and Higashi embeddings were projected to the two-dimensional space with PCA, whereas HiCRep/MDS uses MDS as a dimension reduction method inherently. The embeddings produced by LDA were projected to the two-dimensional space with UMAP as it is the preferred dimension reduction method suggested in Kim et al. (2020).

#### B.2 Simulation data generation for the evaluation of scHi-C imputation

We describe how we generated the simulated scHi-C data for the evaluation of the contact map imputation. For 11,631 imaged chromosome regions (chr21:34.6Mb-37.1Mb) in Bintu et al. (2018), we first turned the 3D coordinates of sequential 30Kb bins into a spatial distance map of size $83 \times 83$ for each cell. As shown in Bintu et al. (2018), the inverse spatial distance between a pair of genomic loci and the number of reads in the corresponding entry of the Hi-C contact map have a high linear correlation. We thus used the inverse spatial distance map as the ground truth (referred to as the probability map) and randomly sampled reads with replacement in proportion to the value of each entry in the probability map. Because the diagonal entries of the distance map are all zeros, which leads to error when trying to calculate the inverse, we directly set all diagonal entries of the probability matrix to zero and excluded the first diagonal for all similarity measurements.

By the law of large numbers, the contact map constructed based on the sampled reads (referred to as sampled contact frequency map) would converge almost surely to the ground truth when the number of sampled reads $n$ goes to infinity. We first evaluated the similarity between the sampled contact frequency map and the ground truth across different read numbers (Fig. S5). As expected, all similarity measurements increase when the read number increases. Moreover, when calculating the distance stratified correlation score (Yang et al., 2017), the correlation scores for long-range interactions are always lower. We observed that only when $n$ is greater than 50,000, the median of both the Pearson and Spearman correlation scores between the simulated contact frequency maps and the ground truth are higher than 0.7, which highlights the significance of contact map imputation. To show the robustness of the method over the sampled read number, we simulated contact maps with 4 groups of read numbers, including 100 to 250, 250 to 500, 500 to 1,000, and 1,000 to 2,500.

We also generated simulated scHi-C datasets based on a more recent 3D genome imaging datasets (3,029 chromosomes, chr2) from Su et al. (2020). Note that this imaging dataset was obtained by labeling and imaging 50Kb segments at intervals of 250Kb. We first approximated the spatial coordinate of a 1Mb segment by averaging the coordinates of the four 50Kb segments within it and then followed the same procedure as above to obtain a spatial distance map of size $243 \times 243$. The probability map was again obtained by calculating the inverse spatial distance map. To reflect the proximity ligation procedure in scHi-C experiments, we set all the entries in the probability map with a distance greater than 2,000 nm in the distance map to zero. The sampled contact frequency maps were then generated based on the probability map. To reflect various coverage of different scHi-C technologies, we calculated the contacts per cell for various scHi-C datasets on chr2 at 1Mb resolution (Table S3) and then chose the read numbers for the simulation data as 250 to 500, 500 to 1,000, 1,000 to 2,500, and 2,500 to 5,000.

In addition, we generated simulated scHi-C datasets by downsampling the WTC-11 scHi-C on chr1 at 1Mb resolution. At a given downsampling rate $1/s$, we randomly selected $1/s$ of all the sequencing reads assigned to a specific cell and generated a sampled contact frequency map. This process was repeated for $s$ times for each cell in the original dataset. We chose the downsampling rate based on the contacts per cell in the existing scHi-C datasets on chr1 at 1Mb resolution (Table S4). The downsampling rates used in this study include 1/3, 1/5, 1/10, and 1/15.

For a fair comparison and also for visualization purpose, all contact maps were quantile-normalized to a standard Gaussian distribution such that the ground truth and the imputation results from different methods would have similar value range and distribution. For the imaging-data based simulation, all the imputation results were evaluated with similarity metrics between the imputed contact map and the ground truth derived from the imaging data. The similarity metrics used include the Pearson correlation score, the Spearman correlation score, and also HiCRep (Yang et al., 2017), which is a stratum-adjusted correlation coefficient score for measuring the similarity of two Hi-C matrices. For downsampling based scHi-C simulation, besides the global Pearson and Spearman correlation scores, we also visualized the distance stratified correlation scores (measured by the Pearson and Spearman for each stratum). Since the full-coverage contact maps (used as the ground truth) are still sparse and noisy, it is sometimes numerically unstable to calculate the correlation score for long-distance interactions. We therefore only included the distance stratified correlation score for the first 15 strata as the full-coverage scHi-C contact maps are still relatively dense in these regions. For the rest of the regions in the contact maps, we calculated the AUPR score with the existence of long-range interactions in the full-coverage contact maps as labels and the values of the corresponding entries in the enhanced contact map as the predicted probability scores.

##### B.3 A/B compartment score annotation for scHi-C dataset

We designed an approach for A/B compartment score annotation based on the widely-used method proposed in Lieberman-Aiden et al. (2009). In the original compartment calling method, the Hi-C contact map is normalized and transformed into a Pearson correlation matrix. The sign of the first principal component (referred to as PC1 for simplicity) of this correlation matrix is then used to define A/B compartments. To generate A/B compartment scores for a scHi-C dataset, the most straightforward way would be to apply the original algorithm to each contact map. However, to compare the A/B compartment annotations across the cell population, this method has several major limitations. First, the original method uses zero as cut-off to binarize the PC1 to A/B compartment label. However, when making comparisons across the cell population, especially within the same cell type, binary labels are not sufficiently expressive to characterize detailed variations. Indeed, subtle changes of compartmentalization that reflects nuclear spatial positioning could lead to important genome function differences (Leemans et al., 2019; Chen et al., 2018; Zhang et al., 2020; Wang et al., 2021). Second, the continuous PC1 scores obtained from individual cells are not directly comparable by definition because the PC1s of different cells are in different lower-dimensional spaces.

To address this, we developed a new method where the first two steps, i.e., normalization and transformation into Pearson correlation matrices, remain the same for each single cell. However, instead of performing PCA on each individual Pearson correlation matrix, we apply PCA once on the Pearson correlation matrix from the pooled scHi-C and save the PCA projection matrix. We then use this bulk projection matrix to transform the single-cell Pearson correlation matrices into continuous one dimensional vectors. Although technically these one dimensional vectors are not the first principal components of each single-cell contact map, for simplicity, we still refer to them as PC1s for each cell. These PC1s in individual cells are continuous and sensitive to reflect the subtle variability of compartment shifts, enabling direct comparison of compartment scores across individual cells.

##### B.4 Calculation of transcriptional variability

In scRNA-seq data, there is an inherent connection between the mean and standard deviation of read count (Hafemeister and Satija, 2019). Therefore, the metrics such as variance or standard deviation would lead to the observation that the highly expressed genes also appear to be the more variable ones. To mitigate this, we quantified the transcriptional variability with two approaches. In the first approach, we use a commonly used metric, coefficient of variation (CV), to balance the correlation between mean and standard deviation (Geiler-Samerotte et al., 2013). Due to the dropout events of the scRNA-seq data, direct calculation of mean and standard deviation will also be less stable on the sparse scRNA-seq data.

We thus utilized MAGIC (Van Dijk et al., 2018) to impute the scRNA-seq data after normalization to address the sparsity issue. The CV is calculated as the ratio between the standard deviation and the mean for each gene on the imputed scRNA-seq data. In the second approach, we used a recently developed method for variance stabilization of scRNA-seq data (Hafemeister and Satija, 2019), which takes into account both the dropout events and the correlation between mean and standard deviation. We applied the method to the scRNA-seq data before imputation. The residual variance is then used as the measurement for the transcriptional variability.

##### B.5 TAD-like domain boundary identification for scHi-C dataset

We used the insulation score based method for calling TAD-like boundaries (Crane et al., 2015). We denote the contact map as $M$ and the window size as $w$. The insulation score $s$ at position $x$ is calculated as:

$$s(x) = \text{inter}(x)/\text{intra}(x) \tag{11}$$

$$\text{inter}(x) = \sum_{x-w\leq i\leq x, x\leq j\leq x+w} M_{ij} \tag{12}$$

$$\text{intra}(x) = \sum_{x-w\leq i\leq x+w, x-w\leq j\leq x+w} M_{ij} \tag{13}$$

A high insulation score indicates that the flanking regions across this genomic loci highly interact with each other, suggesting that this loci is in the middle of a TAD-like domain. On the contrary, a low insulation score indicates that the neighboring loci are well separated and this position is likely a domain boundary. Therefore, the domain boundaries are identified as the local minima of the insulation scores. We used a window size of 500Kb for calculating the insulation scores and identifying the local minima in this work. We kept the window size the same as the original method (Crane et al., 2015).

##### B.6 TAD-like domain boundary calibration for scHi-C dataset

To quantify the cell-to-cell variability of TAD-like domain boundaries in the cell population, it is crucial to have a reliable method for domain boundary comparisons in single cells. Although methods have been developed for comparing a pair or a small number of TAD sets (e.g., Cresswell and Dozmorov (2020); Sauerwald and Kingsford (2018)), it remains computational impractical to directly apply them to scHi-C datasets with hundreds or even thousands of cells due to their poor scalability. We developed a new computational strategy for calibrating domain boundaries called from the scHi-C. Specifically, we first identify a set of domain boundaries shared across the cell population. The size of this shared boundaries is predefined as $K$, which is chosen as 2 times the number of domain boundaries identified in the pooled scHi-C contact maps in this work. During optimization (see below), shared boundaries that are too close to each other ($\leq 1$ bin) would be automatically merged. Each single-cell domain boundary will then be assigned to one of these shared boundaries. In other words, the domain boundaries for each cell will be represented as a subset of this shared boundary set. The assignment and the shared boundary set are optimized such that the calibrated ones are similar to the original ones. Together, this can be defined as:

$$\begin{aligned} \text{minimize}_{B_i^*} \quad & \sum_{i,j} \text{dis}\left(b_{i,j}^*, b_{i,j}\right) \\ \text{s.t.} \quad & b_{i,j} \in B_i \\ & b_{i,j}^* \in B_i^* \\ & B_i^* \subseteq B^* \\ & |B^*| = K, \end{aligned} \tag{14}$$

where $B_i$ and $B_i^*$ represent the single-cell domain boundaries before and after calibration for cell $i$ and $\text{dis}()$ represents the distance measurement of two boundaries. Each shared boundary thus acts as an anchor point such that the variable single-cell domain boundaries associated with it can be analyzed systematically across the cell population. Fig. S10a shows examples of the single-cell domain boundaries before and after calibration. We did not use the domain boundaries called from the pooled scHi-C to specifically allow the algorithm to identify boundaries that are enriched in a group of cells but not strong enough to be captured in the pooled scHi-C. On the other hand, using pooled domain boundaries as initialization for $B^*$ would accelerate convergence. As mentioned, we chose $K$ as 2 times the number of boundaries called from the pooled scHi-C in this work. When initializing $B^*$, half of them are initialized as the pooled domain boundaries while the rest are randomly spaced across the genome.

This problem setting shares similarities with the well-known $K$-means clustering. If we define the $\text{dis}()$ function as the Euclidean distance, it would be equivalent to $K$-means clustering. However, the Euclidean distance based on the genomic coordinate is not appropriate to quantify how well one domain boundary approximates another one (see example in Fig. S10b). When using genomic distance as evaluation metric, both shared domain boundaries have exactly the same distance to the single-cell domain boundary before calibration. However, the left one is preferred as it has relatively similar change in the insulation score. For the similar reason, simply using distance between insulation scores at the domain boundaries is also not desired.

We therefore developed a new metric called "cumulative insulation score" to address this. As illustrated in Fig. S10b, this metric is calculated as the area between the insulation score curve and the horizontal line at the minimum insulation score within the given region. This new metric penalizes the assignment if the assigned domain boundary either has large genomic distance or has very different insulation scores to the original one. With this approach, in both situations mentioned above, the single-cell boundary will be assigned to the correct shared boundary with smaller cumulative insulation score.

Together, we alternate between optimizing the shared boundary set $B^* = \{b_k^*\}$ and optimizing the assignment $c_{i,j}$ until convergence, i.e.,

$$\text{assignment step:} \quad c_{i,j} = \arg\min_k \text{dis}(b_k^*, b_{i,j}) \tag{15}$$

$$\text{update step:} \quad b_k^* = \arg\min_{b_k^*} \sum_{i,j} \mathbb{I}(c_{i,j} = k)\text{dis}(b_k^*, b_{i,j}) \tag{16}$$

where $\text{dis}(b_k^*, b_{i,j})$ is the genomic distance between shared boundary $b_k^*$ and the $j$th boundary in cell $i$. Eventually those shared boundaries with less than 5 assigned single cell boundaries are removed for subsequent analysis.

##### B.7 Identifying differentially expressed genes during WTC-11 differentiation

To probe the connection between variable TAD-like domain boundaries and gene function, we investigated the differentially expressed genes (DEGs) within those boundaries related to differentiation based on scRNA-seq data (Friedman et al., 2018). Specifically, in Friedman et al. (2018), scRNA-seq was acquired from cells at 5 differentiation stages from WTC-11 cells, including pluripotency, germ layer specification, cardiac cells progenitor, committed cardiac cells, and definitive cardiac cells. To filter out low-quality cells, we only kept cells with 500 to 4,000 read counts and less than 10 percent of mitochondrial counts using Seurat (Butler et al., 2018; Stuart et al., 2019). We identified DEGs between pluripotency state and each of the other 4 differentiated stages using Seurat with default parameters.

T. Nagano, Y. Lubling, C. Várnai, C. Dudley, W. Leung, Y. Baran, N. M. Cohen, S. Wingett, P. Fraser, and A. Tanay. Cell-cycle dynamics of chromosomal organization at single-cell resolution. *Nature*, 547

(7661):61, 2017.

V. Ramani, X. Deng, R. Qiu, K. L. Gunderson, F. J. Steemers, C. M. Disteche, W. S. Noble, Z. Duan, and J. Shendure. Massively multiplex single-cell hi-c. *Nature Methods*, 14(3):263, 2017.

N. Sauerwald and C. Kingsford. Quantifying the similarity of topological domains across normal and cancer human cell types. *Bioinformatics*, 34(13):i475–i483, 2018.

T. Stuart, A. Butler, P. Hoffman, C. Hafemeister, E. Papalexi, W. M. Mauck III, Y. Hao, M. Stoeckius, P. Smibert, and R. Satija. Comprehensive integration of single-cell data. *Cell*, 177(7):1888–1902, 2019.

J.-H. Su, P. Zheng, S. S. Kinrot, B. Bintu, and X. Zhuang. Genome-scale imaging of the 3D organization and transcriptional activity of chromatin. *Cell*, 2020.

L. Tan, W. Ma, H. Wu, Y. Zheng, D. Xing, R. Chen, X. Li, N. Daley, K. Deisseroth, and X. S. Xie. Changes in genome architecture and transcriptional dynamics progress independently of sensory experience during post-natal brain development. *Cell*, 2021.

D. Van Dijk, R. Sharma, J. Nainys, K. Yim, P. Kathail, A. J. Carr, C. Burdziak, K. R. Moon, C. L. Chaffer, D. Pattabiraman, et al. Recovering gene interactions from single-cell data using data diffusion. *Cell*, 174(3):716–729, 2018.

Y. Wang, Y. Zhang, R. Zhang, T. van Schaik, L. Zhang, T. Sasaki, D. Peric-Hupkes, Y. Chen, D. M. Gilbert, B. van Steensel, et al. SPIN reveals genome-wide landscape of nuclear compartmentalization. *Genome Biology*, 22(1):1–23, 2021.

T. Yang, F. Zhang, G. G. Yardımcı, F. Song, R. C. Hardison, W. S. Noble, F. Yue, and Q. Li. HiCRep: assessing the reproducibility of Hi-C data using a stratum-adjusted correlation coefficient. *Genome Research*, 27(11):1939–1949, 2017.

L. Zhang, Y. Zhang, Y. Chen, O. Gholamalamdari, Y. Wang, J. Ma, and A. S. Belmont. Tsa-seq reveals a largely "hardwired" genome organization relative to nuclear speckles with small position changes tightly correlated with gene expression changes. *bioRxiv*, page 824433, 2020.

J. Zhou, J. Ma, Y. Chen, C. Cheng, B. Bao, J. Peng, T. J. Sejnowski, J. R. Dixon, and J. R. Ecker. Robust single-cell hi-c clustering by convolution-and random-walk–based imputation. *Proceedings of the National Academy of Sciences*, page 201901423, 2019.

### Supplementary Tables

| Datasets | Cell type / Cell State | Count |
|---|---|---|
| Ramani et al. (Ramani et al., 2017) | GM12878 | 33 |
| | HAP1 | 233 |
| | HeLa | 267 |
| | K562 | 87 |
| | **Total** | 620 |
| 4DN sci-Hi-C (Kim et al., 2020) | GM12878 | 2048 |
| | H1ESC | 2395 |
| | HAP1 | 1338 |
| | HFFc6 | 578 |
| | IMR90 | 29 |
| | **Total** | 6388 |
| Nagano et al. (Nagano et al., 2017) | G1 | 280 |
| | early-S | 303 |
| | late-S/G2 | 326 |
| | mid-S | 262 |
| | **Total** | 1171 |
| WTC-11 | **Total** | 188 |
| sn-m3c-seq. (Lee et al., 2019) | ODC | 1245 |
| | OPC | 203 |
| | Astro | 449 |
| | Endo | 205 |
| | MG | 422 |
| | NN1 | 100 |
| | L2/3 | 551 |
| | L4 | 131 |
| | L5 | 180 |
| | L6 | 86 |
| | Pvalb | 134 |
| | Sst | 217 |
| | Ndnf | 144 |
| | Vip | 171 |
| | **Total** | 4238 |

**Table S1:** The number of cells in all (co-assayed) scHi-C datasets used in this work.

| Datasets | Contacts per cell | | | | Non-zero non-diagonal per cell | | | | Total Possible |
|---|---|---|---|---|---|---|---|---|---|
| | Mean | Min | Median | Max | Mean | Min | Median | Max | |
| Ramani et al. | 10.5k | 2.1k | 7.8k | 109.6k | 4.3k | 1.1k | 3.7k | 25.5k | 237.1k |
| 4DN sci-Hi-C | 5.6k | 1.9k | 3.8k | 752.7k | 4.1k | 1.8k | 3.2k | 107.1k | 223.4k |
| Nagano et al. | 56.9k | 3.1k | 56.8k | 267.1k | 14.7k | 2.5k | 14.1k | 46.5k | 184.5k |
| sn-m3c-seq | 136.6k | 13.5k | 151.3k | 412.9k | 28.0k | 6.0k | 2.9k | 64.1k | 225.0k |
| WTC-11 | 145.1k | 22.5k | 144.8k | 278.3k | 30.6k | 11.6k | 30.5k | 51.5k | 237.1k |

**Table S2:** Statistics of the contacts/non-zero non-diagonal entries per cell at 1Mb resolution for the scHi-C datasets used in this work. The Nagano et al. data were mapped to the mm9 assembly. The Ramani et al. and sn-m3c-seq data were mapped to the hg19 assembly. Others were mapped to the hg38 assembly. Same as Lee et al. (2019), when studying the sn-m3c-seq, only autosomal chromosomes are used.

| Datasets | Contacts per cell | | | | Total Possible |
|---|---|---|---|---|---|
| | Mean | Min | Median | Max | |
| Nagano et al. | 4.4k | 0.2k | 4.4k | 19.4k | 16.5k |
| WTC-11 | 3.3k | 1.2k | 3.3k | 5.2k | 29.4k |
| Ramani et al. | 0.9k | 0.0k | 0.7k | 9.5k | 29.6k |
| 4DN sci-Hi-C | 0.6k | 0.0k | 0.4k | 77.3k | 29.4k |
| sn-m3c-seq | 14.0k | 0.4k | 15.5k | 37.9k | 29.6k |

**Table S3:** Statistics of the contacts per cell at 1Mb resolution of chr2 for different datrasets. The Nagano et al. data were mapped to the mm9 assembly. The Ramani et al. and sn-m3c-seq data were mapped to the hg19 assembly. Others were mapped to the hg38 assembly.

| Datasets | Contacts per cell | | | | Non-zero non-diagonal per cell | | | | Total Possible |
|---|---|---|---|---|---|---|---|---|---|
| | Mean | Min | Median | Max | Mean | Min | Median | Max | |
| Original WTC-11 | 12.1k | 1.2k | 12.1k | 22.9k | 2.9k | 0.9k | 2.9k | 4.4k | 30.9k |
| WTC-11 (1/3) | 4.0k | 0.4k | 4.0k | 7.7k | 1.6k | 0.3k | 1.6k | 2.8k | 30.9k |
| WTC-11 (1/5) | 2.4k | 0.2k | 2.4k | 4.6k | 1.2k | 0.2k | 1.2k | 2.1k | 30.9k |
| WTC-11 (1/10) | 1.2k | 0.1k | 1.2k | 2.4k | 0.8k | 0.1k | 0.8k | 1.4k | 30.9k |
| WTC-11 (1/15) | 0.8k | 0.1k | 0.8k | 1.6k | 0.6k | 0.1k | 0.6k | 1.1k | 30.9k |
| Nagano et al. | 4.5k | 0.2k | 4.6k | 20.4k | 1.2k | 0.2k | 1.2k | 3.8k | 19.5k |
| Ramani et al. | 1.0k | 0.1k | 0.7k | 10.8k | 0.4k | 0.1k | 0.3k | 2.7k | 31.1k |
| 4DN sci-Hi-C | 0.5k | 0.1k | 0.3k | 66.7k | 0.4k | 0.1k | 0.3k | 11.6k | 30.9k |
| sn-m3c-seq | 11.9k | 0.4k | 13.1k | 42.7k | 2.7k | 0.3k | 2.9k | 6.9k | 31.1k |

**Table S4:** Statistics of the contacts/non-zero non-diagonal entries per cell at 1Mb resolution of chr1. The Nagano et al. data were mapped to the mm9 assembly. The Ramani et al. and sn-m3c-seq data were mapped to the hg19 assembly. Others were mapped to the hg38 assembly.

#### Supplementary Figures

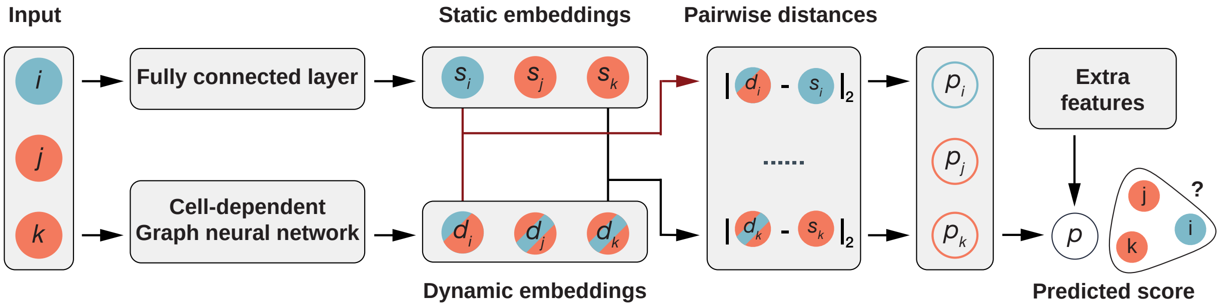


**Figure S1:** The structure of the hypergraph neural network used in Higashi. The input triplet consisting of one cell node and two bin nodes passes through two branches of the network to generate static embeddings and dynamic embeddings for each node, respectively. Then the pairwise distances between static and dynamic embedding pairs are calculated. These pairwise distances are combined with extra features such as genomic distance between the two bins to produce the final predicted score for the input triplet, which represents the probability of an entry in the single-cell contact map.

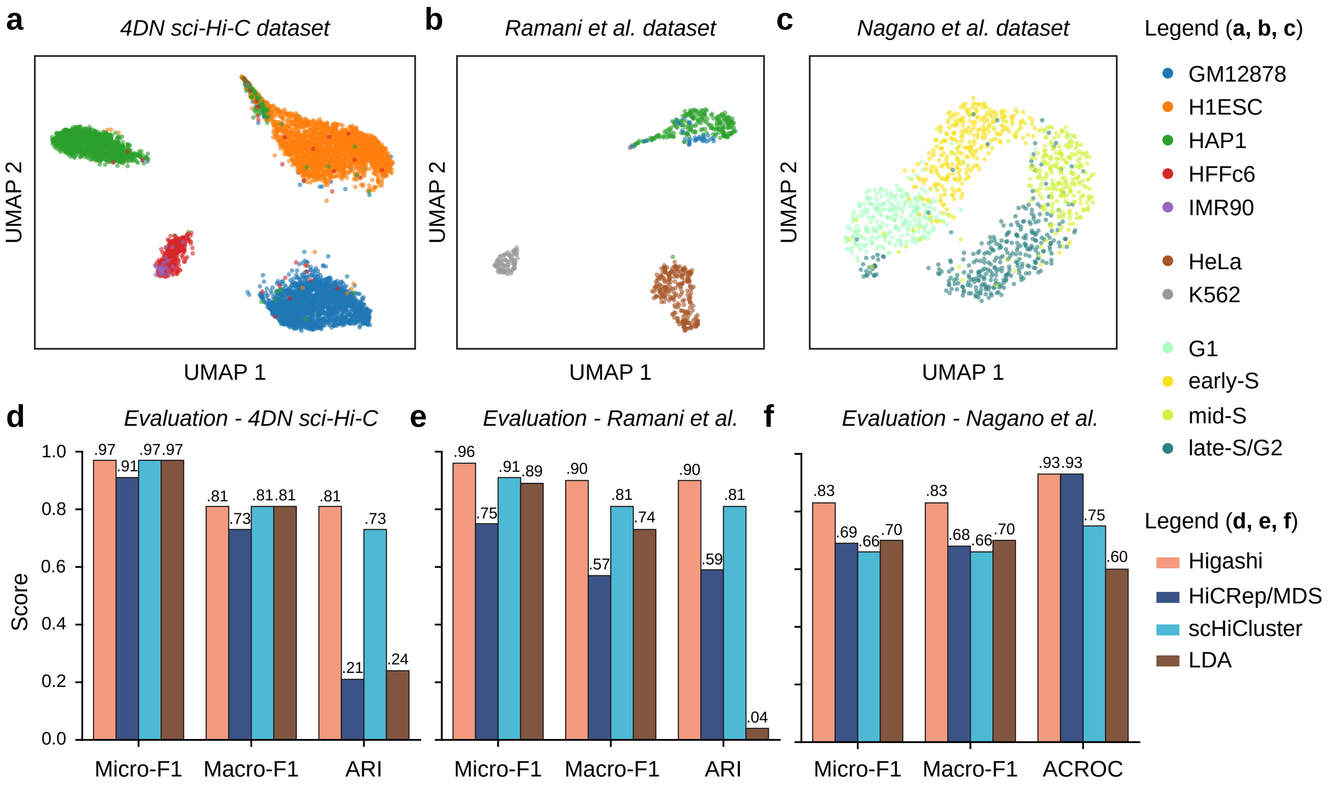


**Figure S2:** Evaluation of the embeddings generated by Higashi. **a, b, c.** The UMAP visualization of the embeddings produced by Higashi for three scHi-C datasets (4DN sci-Hi-C (Kim et al., 2020), Ramani et al. (Ramani et al., 2017), and Nagano et al. (Nagano et al., 2017) datasets). **d, e, f.** Quantitative evaluation of Higashi on the three scHi-C datasets by comparing to HiCRep/MDS (Liu et al., 2018), scHiCluster (Zhou et al., 2019), and LDA (Kim et al., 2020). The performances are measured by Micro-F1, Macro-F1, Adjusted Rand Index (ARI), and also ACROC scores from the supervised and unsupervised cell type identification tasks.

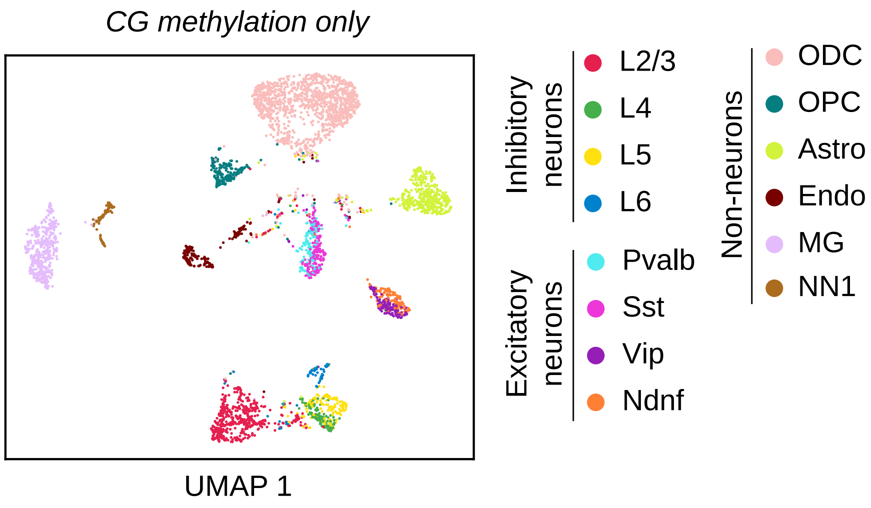


**Figure S3:** UMAP visualization of the embeddings using CG methylation only. The embeddings are generated by Scanorama (Hie et al., 2019). The dimension size is chosen as 64.

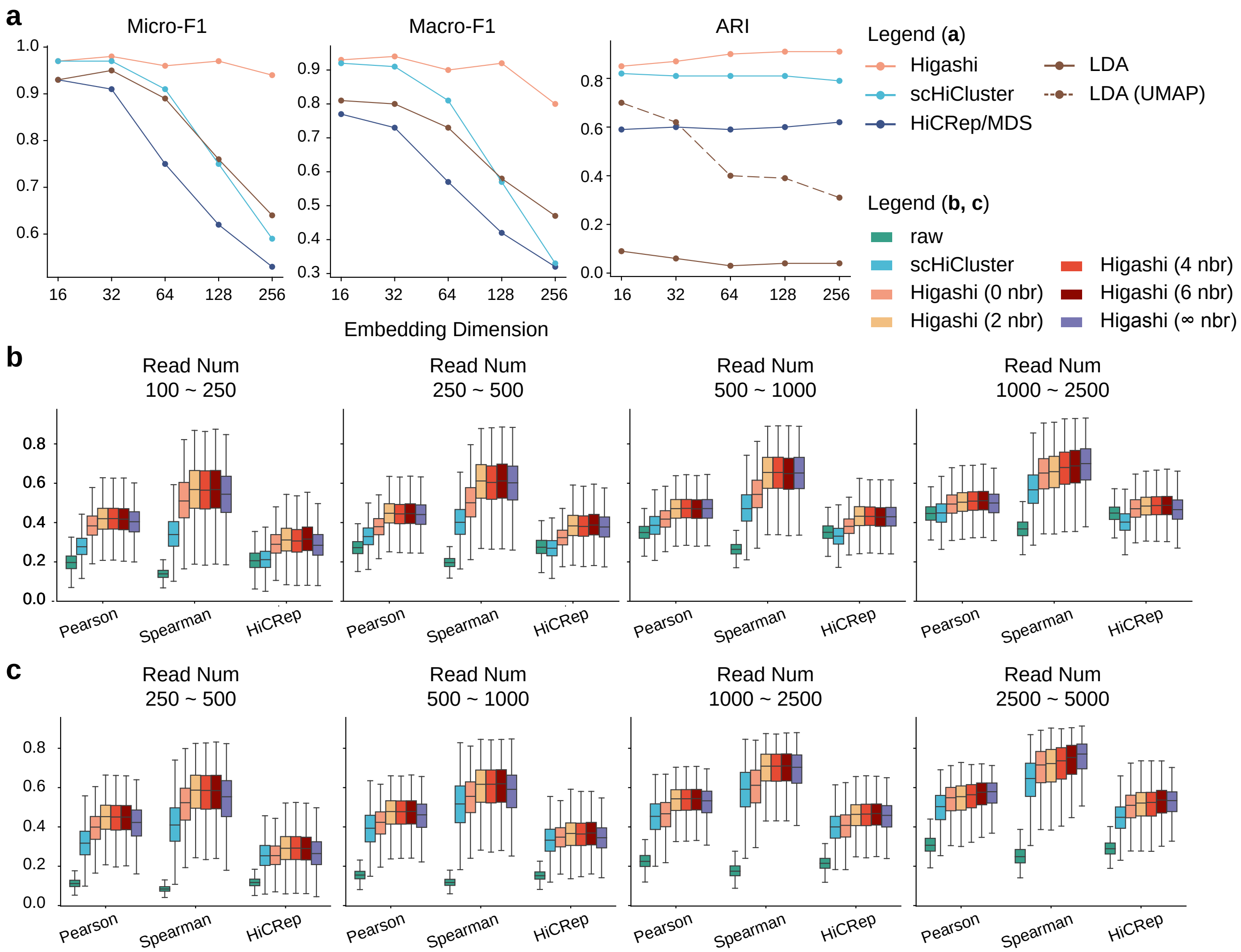


**Figure S4:** Higashi are robust to hyperparameter selections. **a.** The performance of the supervised and unsupervised cell type identification tasks measured by Micro-F1, Macro-F1, and Adjusted Rand Index (ARI) scores over different scHi-C analysis methods (Higashi, HiCRep/MDS (Liu et al., 2018), scHiCluster (Zhou et al., 2019) and LDA (Kim et al., 2020)) and different embedding dimensions. The experiment was done on the Ramani et al. dataset. **b, c.** The performance of the contact map imputation measured by the Pearson correlation, Spearman correlation, and HiCRep (Yang et al., 2017) scores over the baseline methods and Higashi with different number of neighboring cells in the embedding space for imputation. The evaluation in **(b)** was done using the simulation data based on the imaging data from Bintu et al. (2018). The experiment in **(c)** was done using the simulation data based on the imaging data from Su et al. (2020).

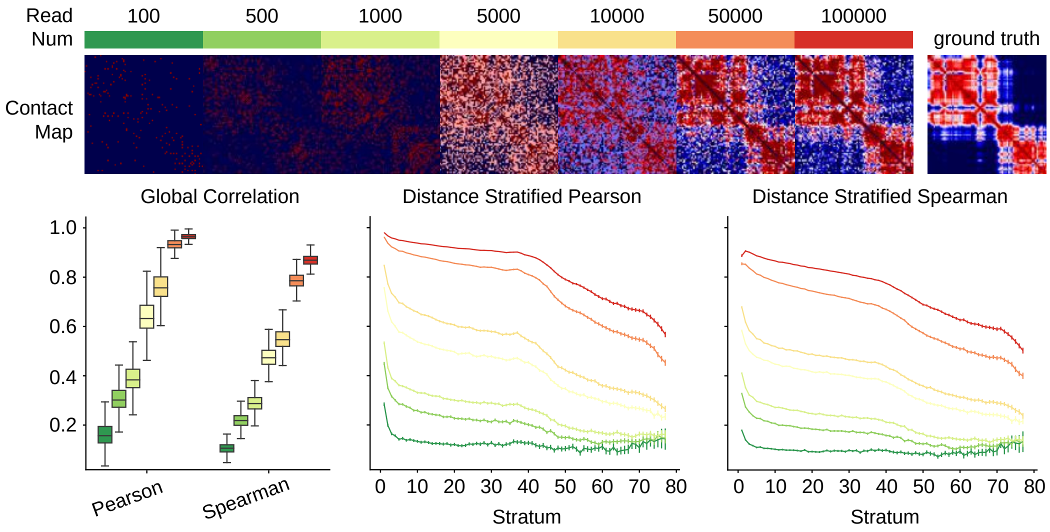


**Figure S5:** Similarities between the simulated contact frequency maps and the ground truth probability map over different read numbers. Simulation is based on the 3D chromosome imaging data from Bintu et al. (2018). The similarities are quantified by the global Pearson and Spearman correlation scores as well as the distance stratified Pearson and Spearman correlation scores.

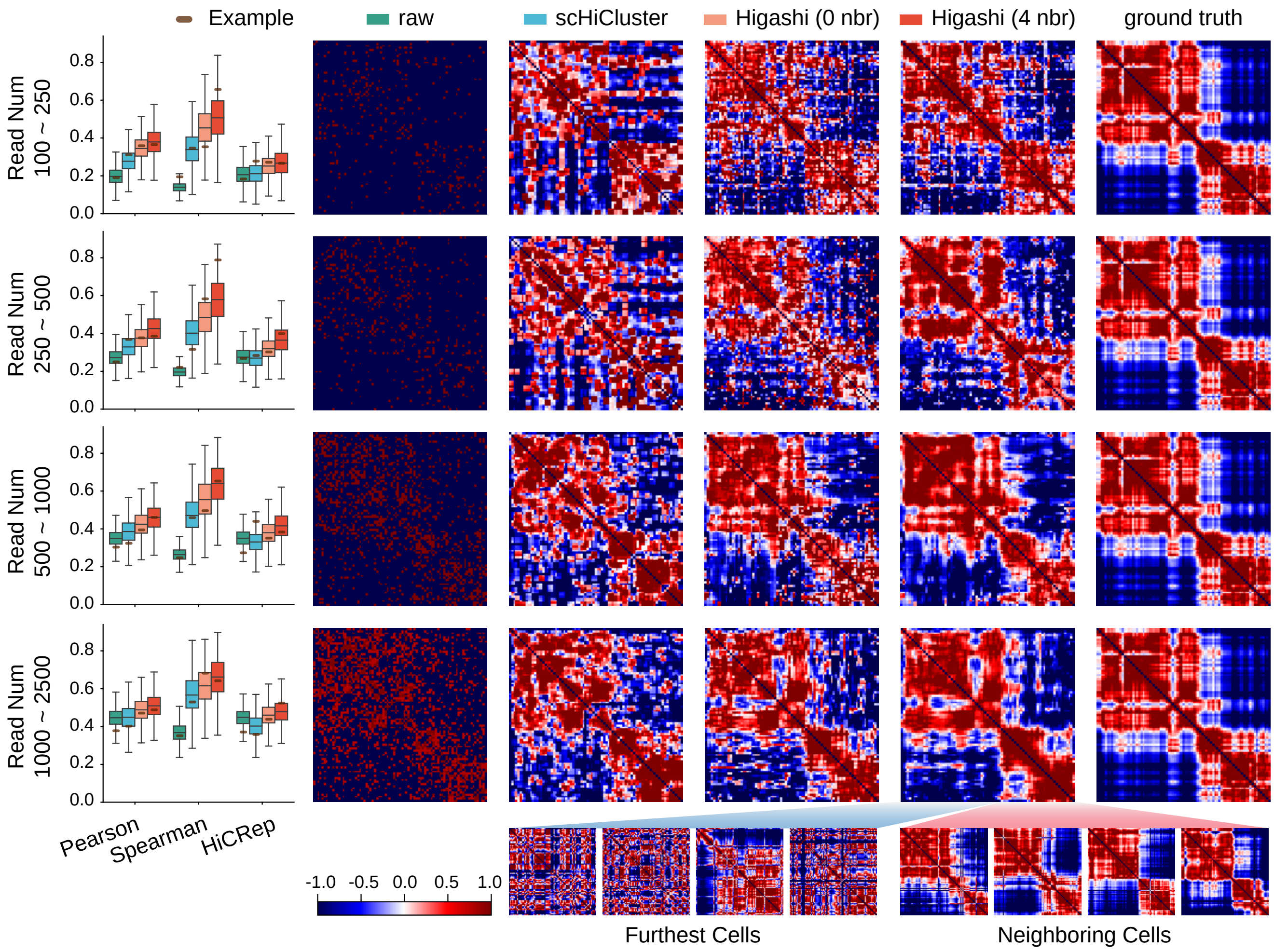


**Figure S6:** Evaluation and visualization of different imputation methods on the simulated scHi-C data based on the 3D chromosome imaging data from Bintu et al. (2018). Each row corresponds to one set of simulation data with a chosen range of the read number. The boxplots illustrate the quantitative evaluation of the similarities between the raw, the scHiCluster enhanced, the Higashi enhanced contact maps, and the ground truth (inverse distance map). The heatmaps visualize the contact map before and after imputation as well as the ground truth. The contact maps of the neighboring cells in the embedding space that help the imputation are included. As a reference, the contact maps of the cells that are furthest in the embedding space are also visualized. Related to Fig. 2e.

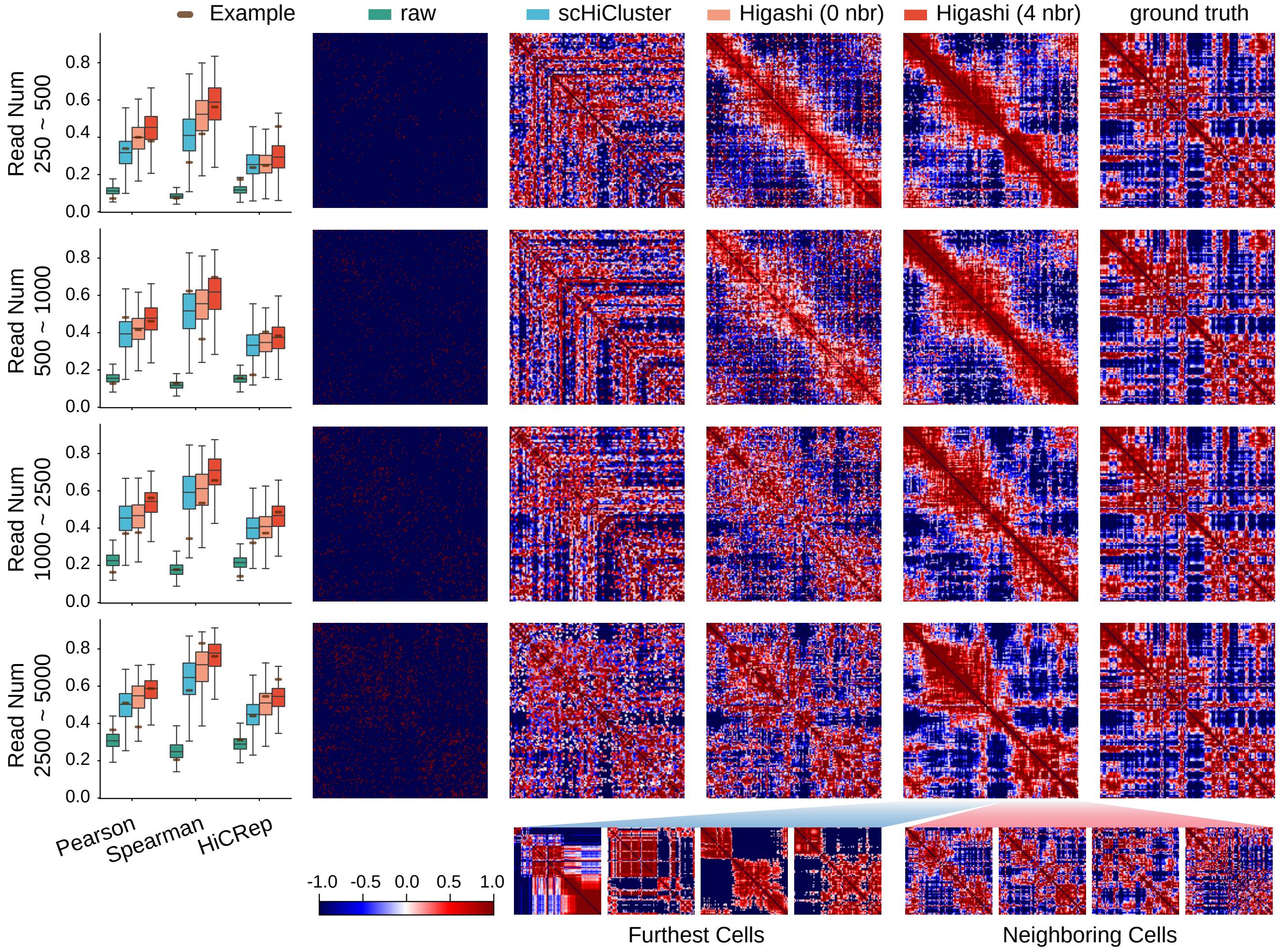


**Figure S7:** Evaluation and visualization of different imputation methods on the simulated scHi-C data based on the 3D genome imaging data from Su et al. (2020). Each row corresponds to one set of simulation data with a chosen range of the read number. The boxplots illustrate the quantitative evaluation of the similarities between the raw, the scHiCluster enhanced, the Higashi enhanced contact maps, and the ground truth (inverse distance map). The heatmaps visualize the contact map before and after imputation as well as the ground truth.

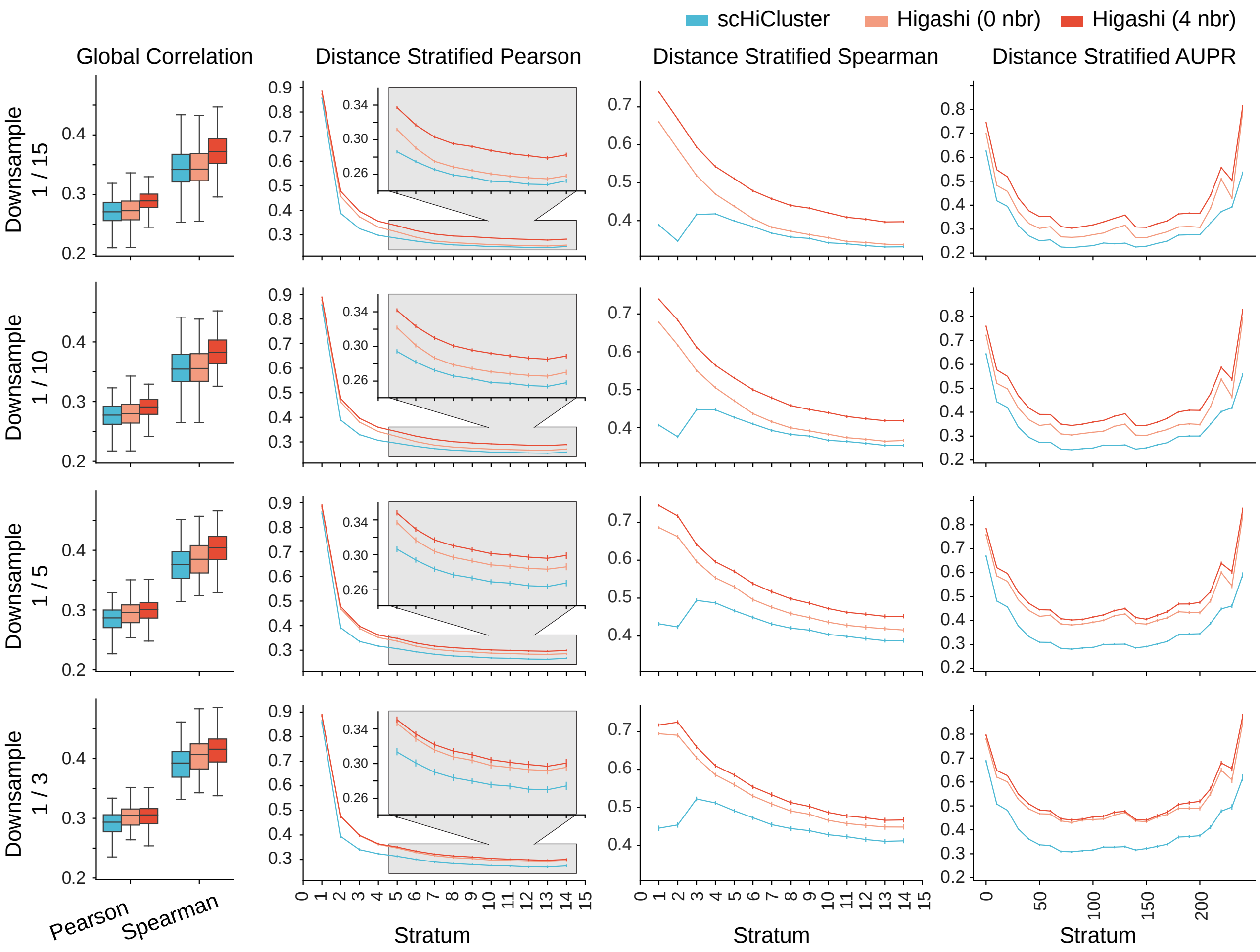


**Figure S8:** Evaluation of different imputation methods on the scHi-C data simulated by downsampling the WTC-11 scHi-C datasets. Each row corresponds to one set of downsampling rate. The boxplots illustrate the global correlation scores of contact maps before and after imputation versus the ground truth. The line plots visualize the evaluation metrics versus each stratum. The evaluation metrics include the Pearson correlation and Spearman correlation for short-range interactions as well as AUPR (area under precision-recall) score for long-range interactions.

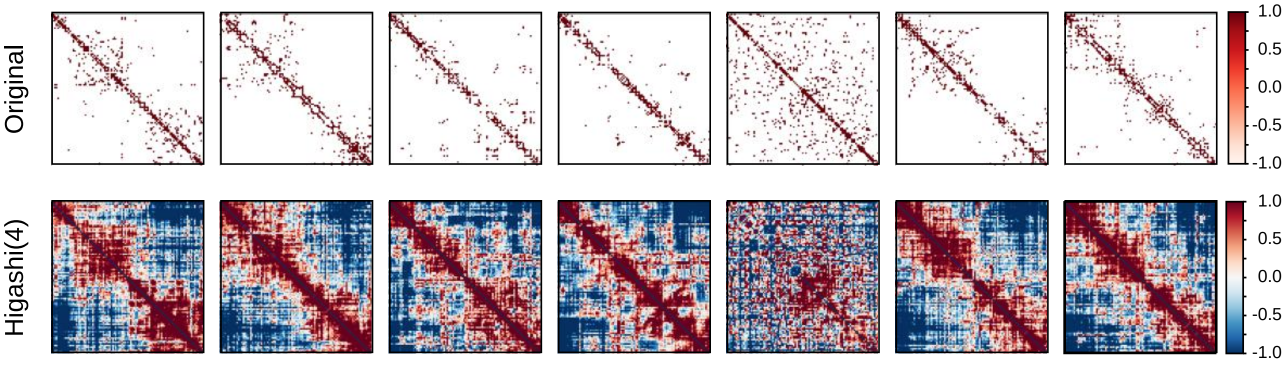


**Figure S9:** Examples of the Higashi imputation on WTC-11 scHi-C at 50Kb resolution (chr21:15Mb-20Mb)

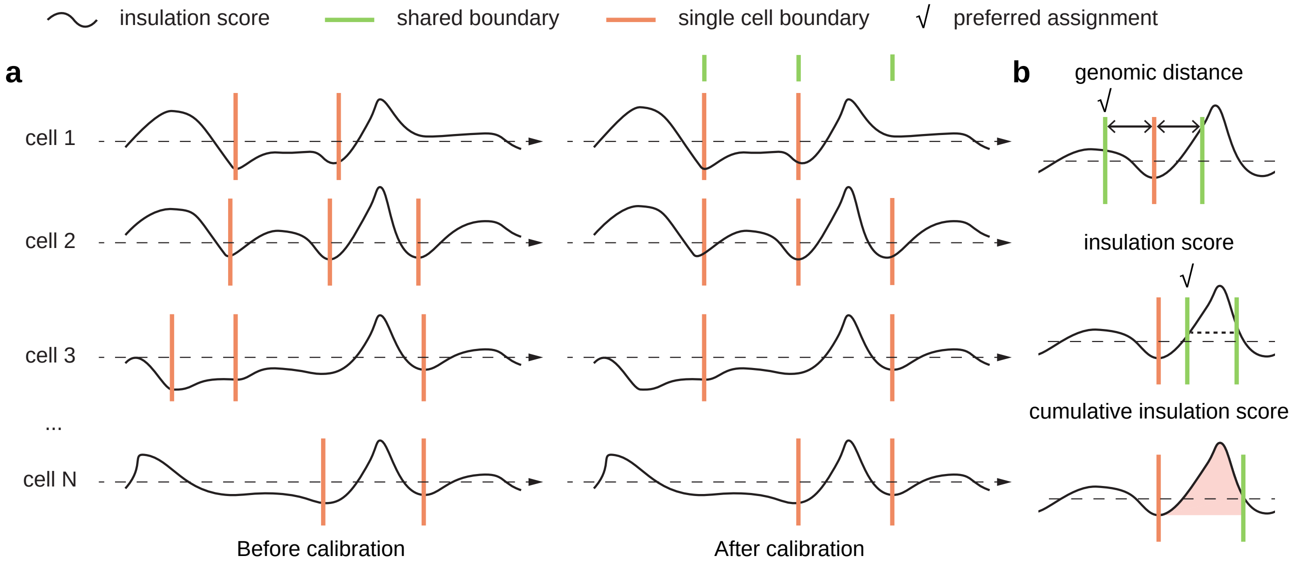


**Figure S10:** Illustration of TAD-like domain boundary calibration among single cells. **a.** Domain boundary calibration for boundaries called from scHi-C. **b.** Different measurements for comparing similarities of two domain boundaries with the given insulation score. Cases where the given metric cannot differentiate the preferred boundaries from the less appropriate ones are also visualized.

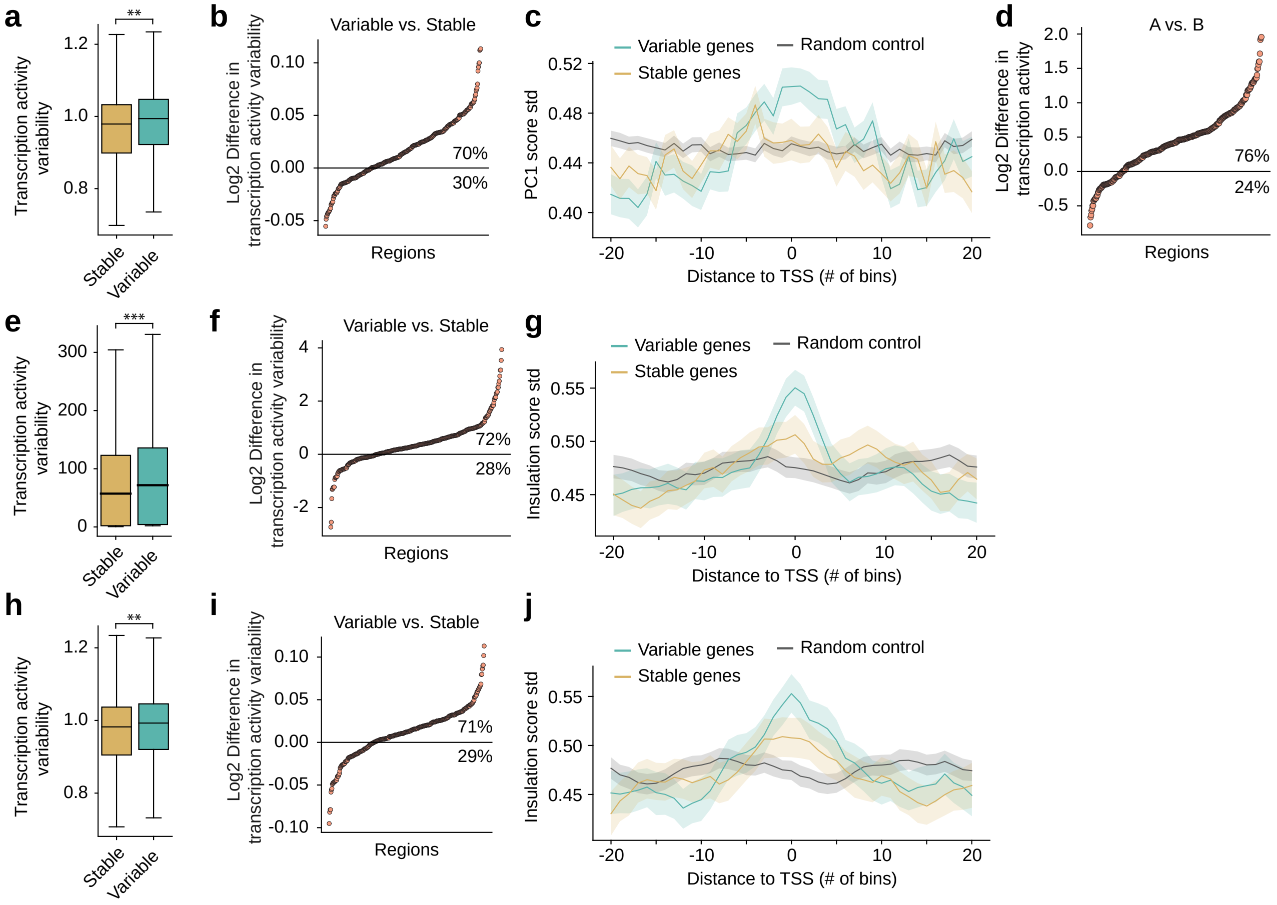


**Figure S11:** Correlation between the cell-to-cell variability of 3D genome features (A/B compartment scores and TAD-like domain boundaries) and transcriptional variability. **a.** Global comparisons of transcriptional variability on regions with variable and stable compartment annotations. There are 3,900 genes that have stable single-cell compartment scores with average transcription activity variability equal to 0.97. There are 3,890 genes that have dynamic single-cell compartment score with average transcription activity variability equal to 0.96. The one sided t-test *P*-value = $9.0 \times 10^{-3}$ **b.** Log2 difference of transcriptional variability of genes with variable versus stable compartment annotations within Mb scale windows. **c.** Visualization of standard deviation of the compartment scores around genes with variable or stable transcription level. **d.** Log2 difference of the average transcription of genes in A compartments versus genes in B compartments within Mb scale windows. **e, h.** Global comparisons of transcriptional variability on regions with variable and stable insulation scores. In **(e)**, there are 5,075 genes that have stable single-cell compartment scores with average transcription activity variability equal to 86.4. There are 5,071 genes that have dynamic single-cell compartment score with average transcription activity variability equal to 77.1. The one sided t-test *P*-value = $1.1 \times 10^{-8}$. In **(h)**, here are 3,900 genes that have stable single-cell compartment scores with average transcription activity variability equal to 0.97. There are 3,890 genes that have dynamic single-cell compartment score with average transcription activity variability equal to 0.96. The one sided t-test *P*-value = $9.0 \times 10^{-3}$. **f, i.** Log2 difference of transcriptional variability of genes with variable versus stable insulation scores within a Mb scale window. **g, j.** Visualization of standard deviation of insulation scores around genes with variable or stable transcription level. In panels **a,b,c,h,i,j**, the transcriptional variability is quantified as the residual variance calculated by a variance stabilization method developed for scRNA-seq data (Hafemeister and Satija, 2019). In panels **e,f,g**, the transcriptional variability is quantified as the coefficient of variation of the imputed scRNA-seq data (Van Dijk et al., 2018). Related to Fig. 3b,c,d

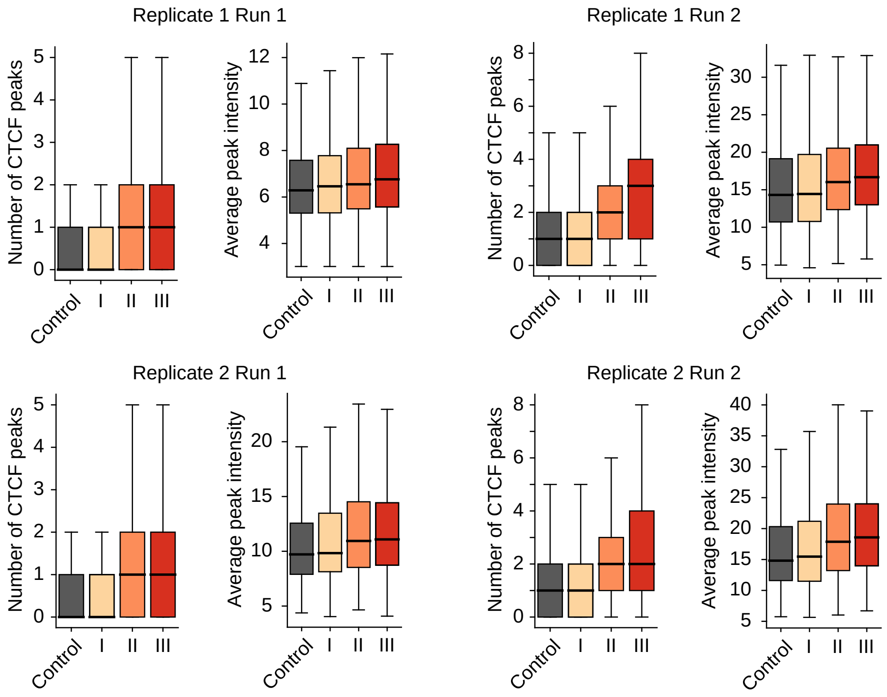


**Figure S12:** CTCF binding at domain boundaries from different occurrence frequency groups across the cell population. CTCF binding peaks were obtained on each batch from each replicate individually. Related to Fig. 3i,j.

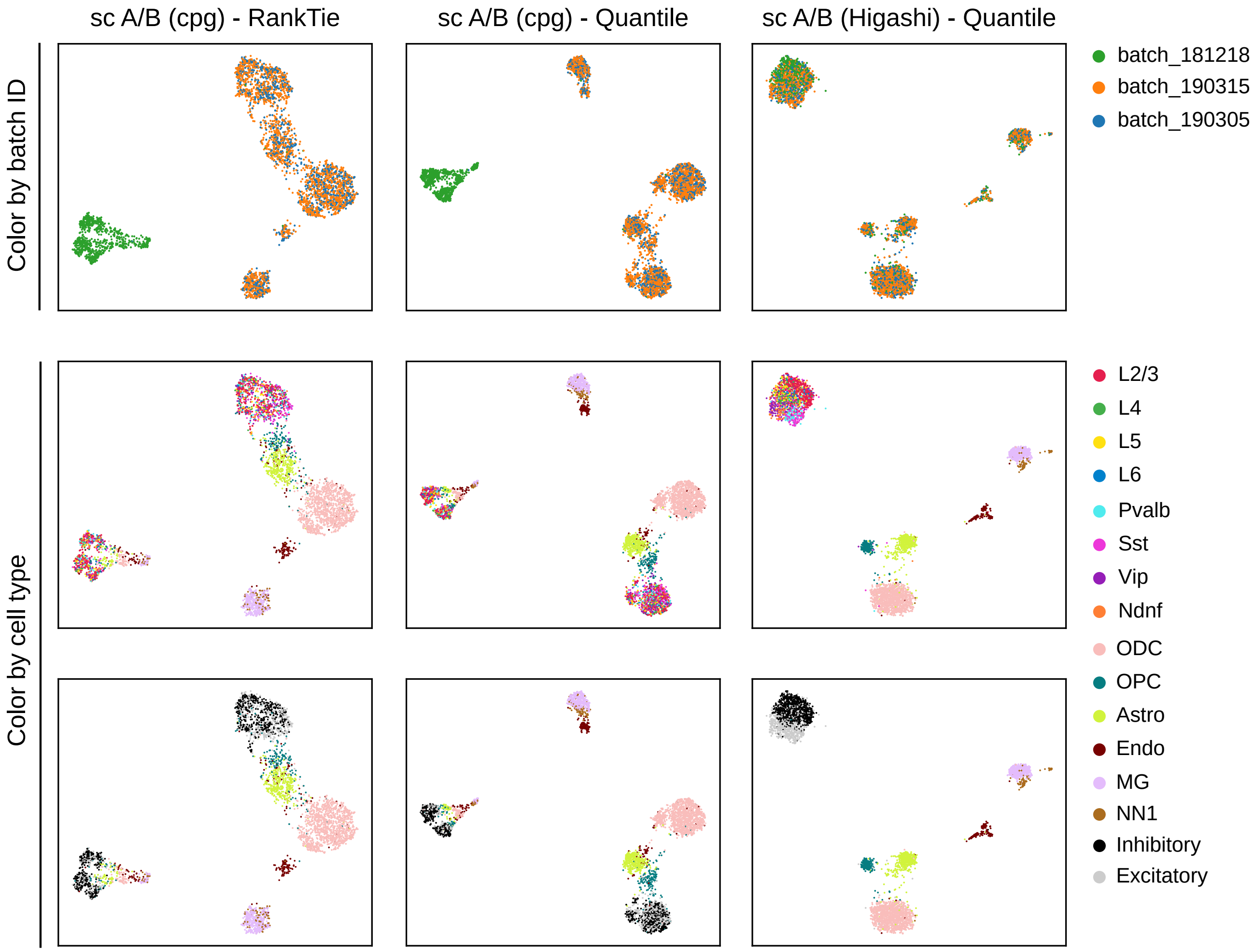


**Figure S13:** UMAP visualization of the single-cell A/B compartment values. Each row corresponds to one color scheme while each column corresponds to a specific combination of calculation method and normalization method for single-cell A/B compartment scores. The calculation method includes the approach using CpG density as approximation (as used in Tan et al. (2021)) and the method proposed in this work. The A/B compartment values are further normalized with either RankTie (as used in Tan et al. (2021)) or quantile transformation.

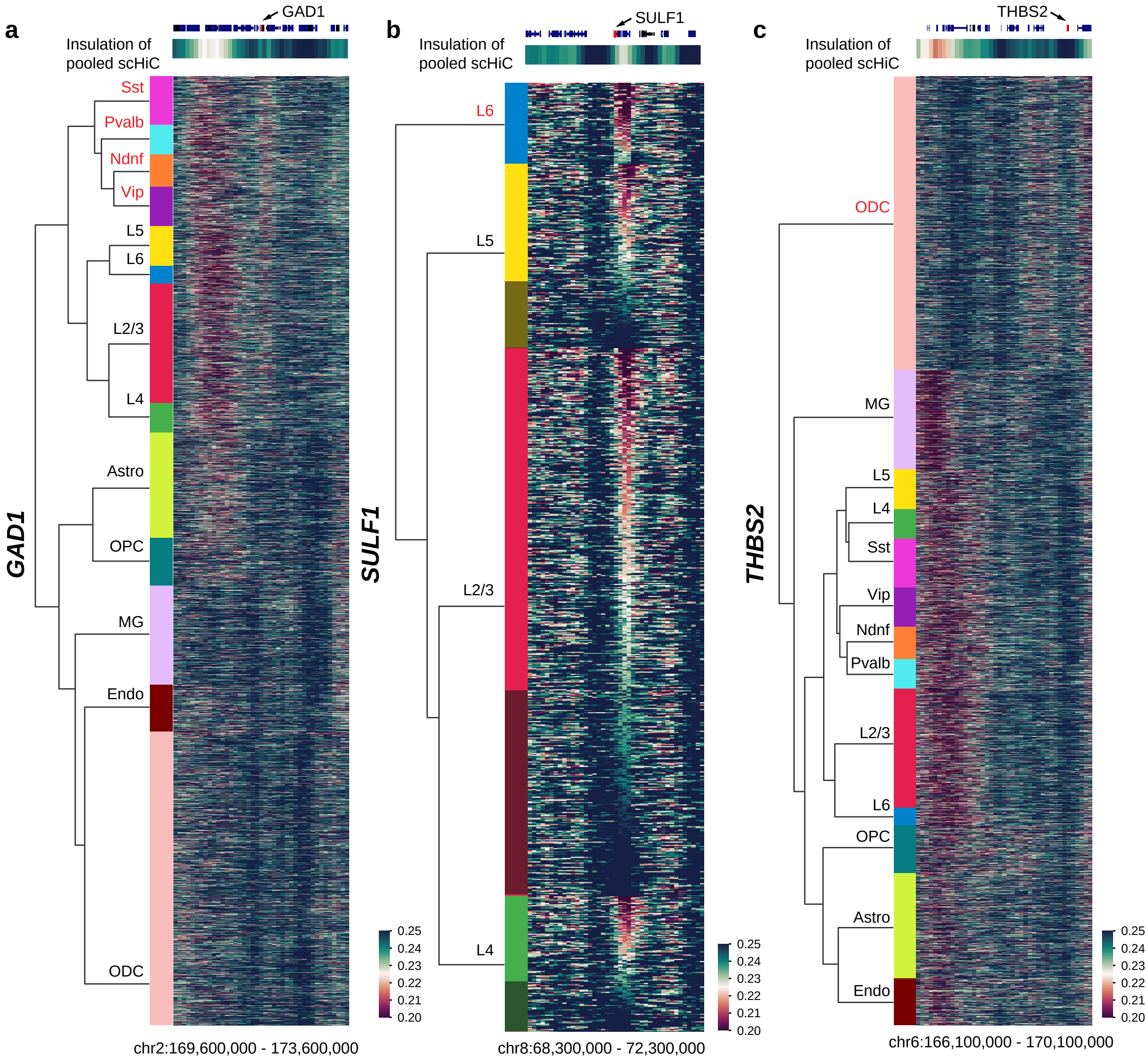


**Figure S14:** Single-cell insulation score heatmap of the flanking regions surrounding the genes in different cell types with distinct TAD-like boundary patterns specific to the highlighted cell type (red rectangle). **a.** Marker gene GAD1 for inhibitory neuron subtypes Sst, Pvalb, Ndnf, and Vip. **b.** Marker gene SULF1 to distinguish L6 subtypes from the rest excitatory neuron subtypes (L2/3, L4, L5). Cells within each cell type are sorted according to the average insulation scores around the TAD-like domain boundary of interest. Cells without the presence of such boundary are shaded. **c.** THBS2 for ODC. THBS2 has ODC-specific high expression. Insulation scores that are calculated based on the pooled scHi-C contact maps of the whole population are also visualized as a reference at the top.

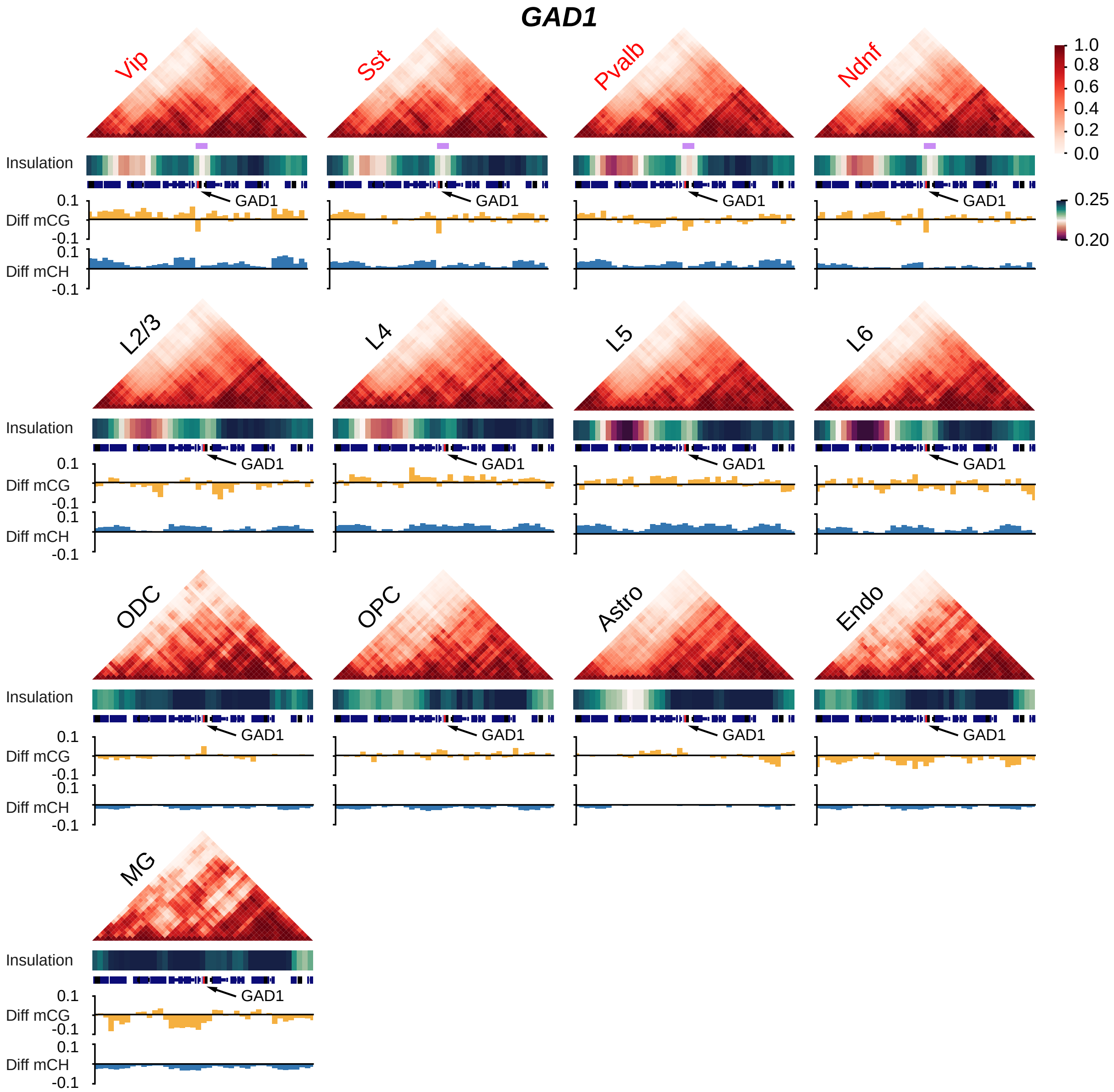


**Figure S15:** Pooled imputed contact maps, insulation scores, and methylation profiles of the flanking regions (+/- 2Mbp) of the marker gene GAD1 for inhibitory neuron subtypes Sst, Pvalb, Ndnf, and Vip. Light purple bar shows a TAD-like domain boundary specific to inhibitory neuron subtypes. Related to Fig. 4d.

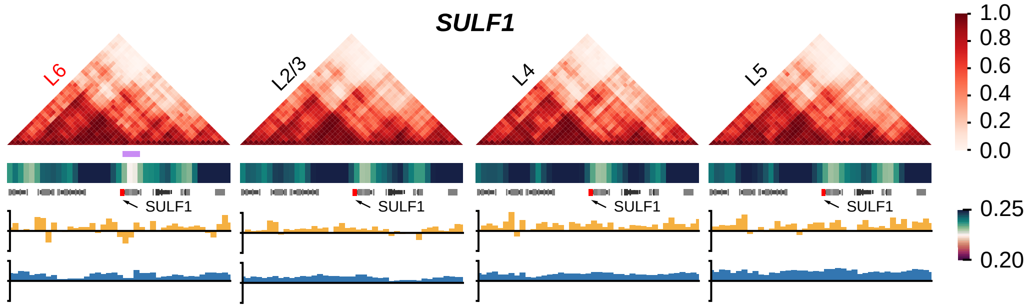


**Figure S16:** Pooled imputed contact maps, insulation scores, and methylation profiles of the flanking regions (+/- 2Mbp) of the marker gene SULF1 to distinguish L6 subtypes from the rest excitatory neuron subtypes (L2/3, L4, L5). Light purple bar shows a stronger TAD-like domain boundary specific to L6.

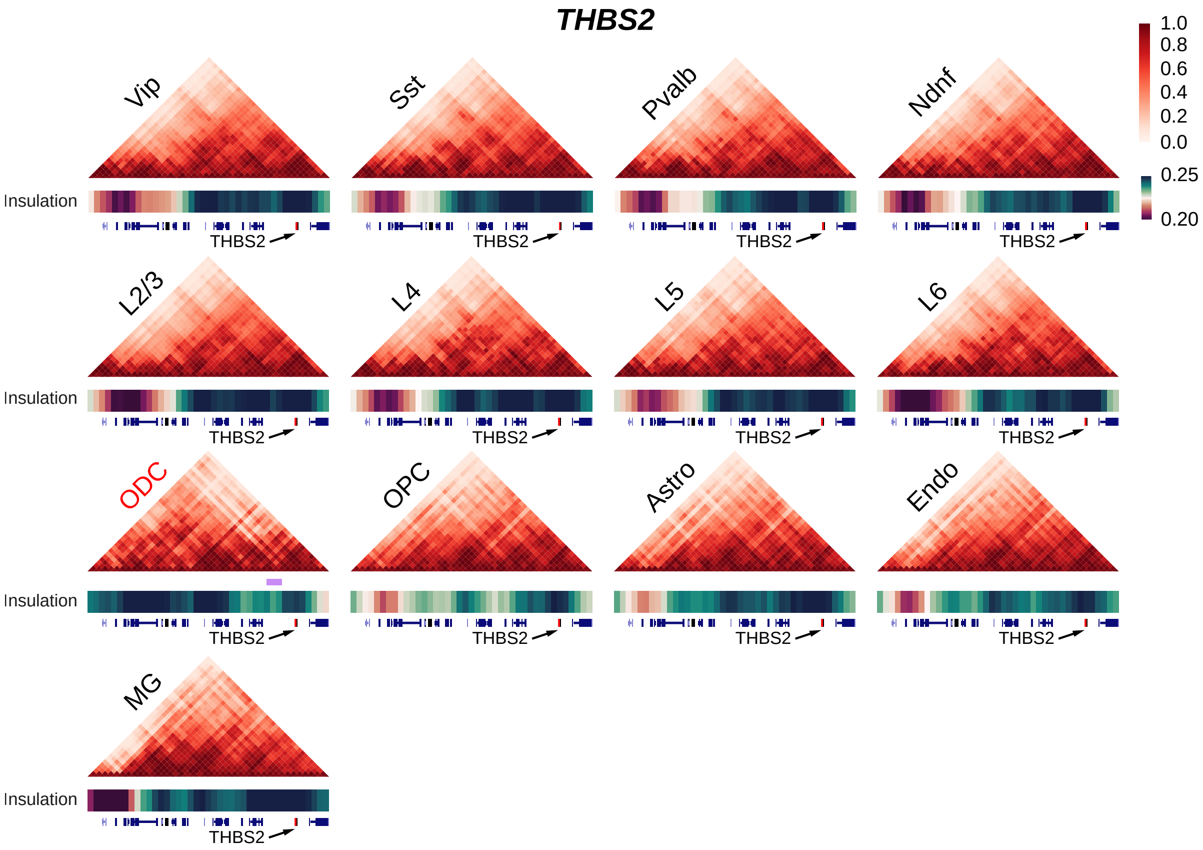


**Figure S17:** Pooled imputed contact maps, insulation scores, and methylation profiles of the surrounding regions (+0.5Mbp/-3.5Mbp) of the gene THBS2 that is near a cell-type TAD-like boundary in ODC and has cell-type specific high expression in ODC. Light purple bar shows an ODC-specific TAD-like domain boundary. Related to Fig. 4f.

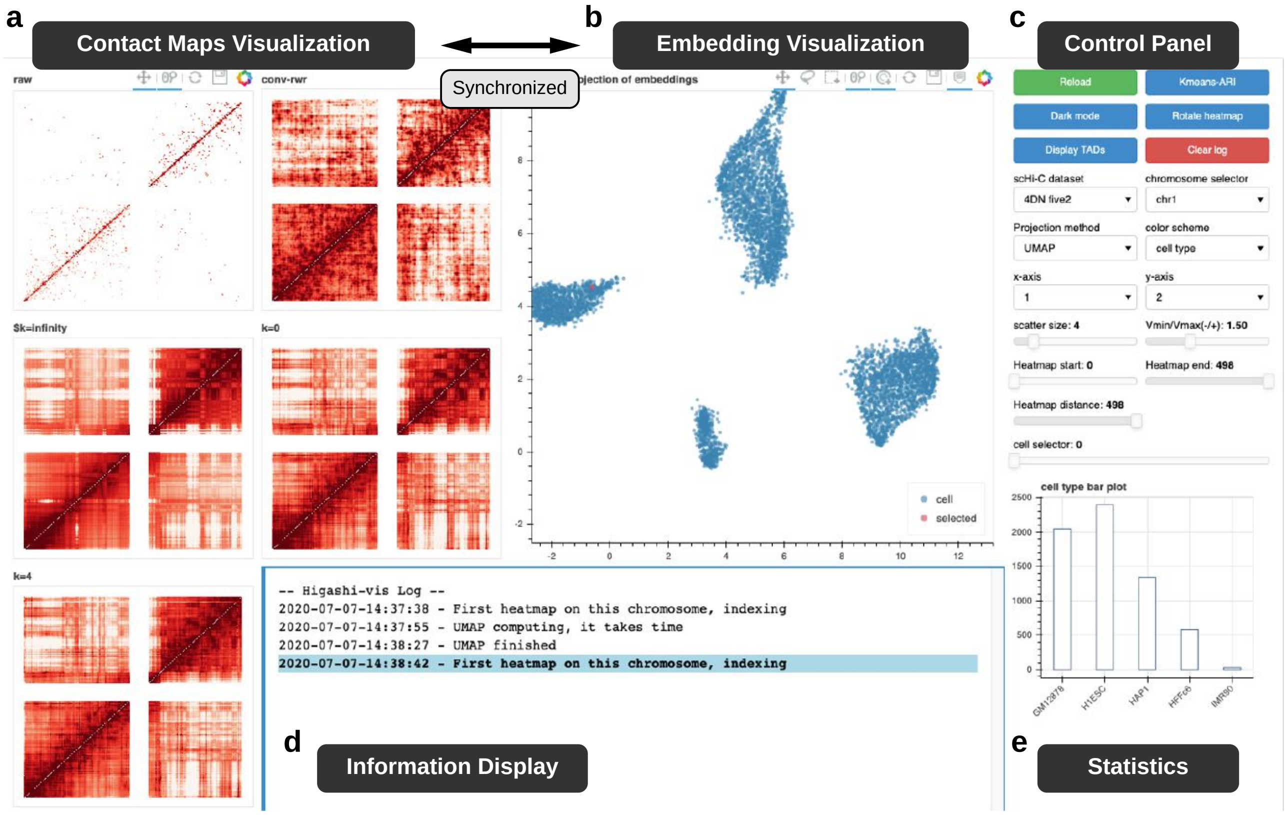


**Figure S18:** Screenshot of the visualization tool in Higashi. **a.** Visualization of the single cell (pooled) contact maps before and after imputation with different methods. This panel is synchronized for visualization with panel **b**. **b.** Visualization of the scHi-C embeddings where each dot corresponds to one cell. Clicking or selecting cell(s) of interest will update the panel **a** interactively. **c.** Control panel for the visualization such as the color scheme of the embedding scatter plot in panel **a,b**. **d.** Display messages from the server. **e.** Visualization of statistics of the cell population as well as the selected cells in panel **b**. Note that this panel will also update interactively.
